# The baculovirus promoter OpIE2 sequence has inhibitory effect on the activity of the Cytomegalovirus (CMV) promoter in HeLa and HEK-293T cells

**DOI:** 10.1101/2020.10.09.315515

**Authors:** A Aladdin, N Sahly, R Faty, MM Youssef, TZ Salem

## Abstract

Understanding how promoters work in non-host cells is complex. Nonetheless, understanding this process is crucial while performing gene expression modulation studies. In this study, inhibitory regions in the 5’ end of the OpIE2 insect viral promoter were found to be blocking the activity of the CMV promoter in mammalian cells. This finding was reached in the process of constructing a shuttle vector with CMV and OpIE2 promoters in a tandem arrangement to achieve gene expression in both mammalian and insect cells, respectively. OpIE2 promoter was cloned downstream of the CMV promoter and upstream of the EGFP reporter gene. After introducing the constructed shuttle vector to insect and mammalian cells, a significant drop in the CMV promoter activity in mammalian cells was observed. To enhance the CMV promoter activity, several modification were made to the shuttle vector including site-directed mutagenesis to remove all ATG codons from the downstream promoter (OpIE2), separating the two promoters to eliminate the effect of transcription interference between them, and finally, identifying some inhibitory regions in the OpIE2 promoter sequence. When these inhibitory regions were removed, high expression levels in insect and mammalian cells were restored. In conclusion, a shuttle vector was constructed that works efficiently in both mammalian and insect cell lines. This study showed that inserting 261 to 313 bp from the 3’ end of the OpIE2 promoter downstream of the CMV promoter maintains efficient gene expression in both Sf9 and mammalian cells.

## INTRODUCTION

The use of shuttle vectors has become a time and cost-efficient solution to clone target genes that require expression in different species. However, to allow any gene of interest to be expressed in different species with only one cloning step, the promoters need to be arranged in tandem.

Most shuttle vectors are designed to work in multiple bacterial species or in both bacterial and eukaryotic hosts but to a lesser extent in two different eukaryotic species. One of the reasons is that bacterial promoters are usually shorter than the eukaryotic promoters, therefore, combining them is easier than combining two different eukaryotic promoters that are usually long. Shuttle vectors that can drive constitutive gene expression in both mammalian and insect cells are uncommon. Early studies (Koedood et al. 1995; Tomita et al. 1995) combined two strong promoters working in mammalian and insect cells, cytomegalovirus (CMV) immediate early promoter and the baculovirus p10 promoter, respectively. The p10 promoter is relatively short (∼121 bp) and only works during baculovirus infection that eventually leads to cellular lysis (Tomita et al. 1995). In that study, the CMV promoter (first in order) was placed upstream of the p10 promoter (second in order) to avoid having any ATGs in the second promoter. This arrangement also ensures the proper distance between the first promoter and the target genes, as the p10 promoter is around five times shorter than the CMV promoter (Philipps et al. 2008). Morevoer, there is no known distance dependence for the CMV promoter (Kozak 1999).

The immediate early OpIE2 promoter from the baculovirus Orgyia pseudotsugata multicapsid nucleopolyhedrovirus (OpMNPV) can serve to circumvent the cellular lysis process during baculovirus infection (Bleckmann et al. 2015). It is a constitutive promoter that does not require baculovirus infection to work and can induce gene expression in various insect cell lines such as Hi5, Sf9, Sf21, and LD652Y (Theilmann and Stewart 1992). OpIE2 promoter was found to have the highest activity in both Hi5 and Sf21 insect cell lines when compared to other early viral promoters such as IE1, hr5IE1p10 (a cassette of hr5, E1, and p10), or other endogenous Sf21 insect promoters (Bleckmann et al. 2015). The OpIE2 promoter alone was found to be inactive in many mammalian cell lines, and similarly the CMV promoter was inactive in insect cells (Pfeifer et al. 1997; Cheng 2004).

In the present study, we test the possibility of maintaining high activity of the commonly used constitutive promoters, the CMV promoter in mammalian cells and OpIE2 promoter in insect cells, when placed in tandem. OpIE2 promoter was placed downstream of the CMV promoter and upstream of the reporter gene (EGFP). While OpIE2 promoter activity should not be affected in insect cells being closer to the target gene, the activity of the first promoter (CMV) was compromised in mammalian cells. Several factors were studied to understand the reason behind this low CMV promoter activity in mammalian cells, including the presence of ATG trinucleotides in the OpIE2 promoter sequence and transcriptional interference. After investigating these possibilities, we hypothesized the presence of inhibitory sequence(s) within the OpIE2 promoter that compromised CMV promoter activity on the transcription level. Several regions were identified at the 5’ end of the OpIE2 promoter sequence that inhibited CMV promoter activity in mammalian cells. Removing these inhibitory regions from the OpIE2 promoter sequence retained the majority of OpIE2 and CMV promoters’ activities in insect and mammalian cells, respectively. Further 5’ end deletions of the OpIE2 promoter sequence beyond −261 bp from the transcription start site gradually enhanced the CMV promoter activity but reduced the OpIE2 promoter activity in Sf9 insect cells.

## MATERIALS AND METHODS

### *In silico* Analysis of OpIE2 Promoter Sequence

The sequence of the OpIE2 promoter was obtained from pIB/v5-His map sequence (ThermoFisher Scientific, USA). The truncated version of the wild-type OpIE2 promoter sequence (tIE2), site-directed mutagenesis (SDM) versions of the tIE2, and a sequence of a chimeric fusion of CMV promoter and tIE2 promoter (CMV/tIE2) were analyzed on TRANSFAC database version 2017.3 (Matys 2003) using Match analysis tool (Kel 2003) that utilizes insects’ transcription factors library. This analysis was done to see the effect of fusing the two promoters on the transcription binding pattern of the OpIE2 promoter in insect cells. The database relies on position weight matrices (PWMs), based on insect transcription factors’ profile. The obtained data were used to build a profile for the putative transcription factors and their binding sites.

### Clones Construction and Site-Directed Mutagenesis

The full-length OpIE2 promoter was obtained from the pIB/V5-His by PCR using *Phusion* High fidelity DNA Taq polymerase (ThermoFisher Scientific, USA) according to the following program: 98°C for 30 s followed by 30 cycles of (98°C for 30 s, 60°C for 30 s, 72°C for 2 min) and a final extension step at 72°C for 10 min. The PCR product was amplified by the forward primer: *OpIE2/F1 5’-TAT<u>CTCGAG</u>GGATCATGATGATAAACAAT-3*’ and the reverse primer: *OpIE2/R1 5’-TA<u>GAATTC</u>TAAATTCGAACAGATGCTG-3’*; before cloning into pEGFP-N1 vector (Clontech, USA) between *Xho*I and *Eco*RI sites, to obtain the chimeric CMV/OpIE2 promoter sequence upstream of the *EGFP* gene. The truncated version of OpIE2 (tIE2) was created by deleting 70 bp from the 5’ end of the wild-type promoter sequence using the forward primer OpIE2/F2: *5’-TAT<u>CTCGAG</u>TTTGCCAACAAGCACCTT-3’* and the reverse primer (OpIE2/R1). ATG-free versions of the truncated OpIE2 promoter sequence (tIE2 promoter with no ATG triplet in its sequence) were created by Q5 Site-Directed Mutagenesis (SDM) Kit (New England Biolabs-NEB, USA). Primers used to generate mutations were designed using the NEBaseChanger tool (NEB, USA), listed in Table S1. All mutated clones (pGTG, pCTG, pTTG, pAAG, pACG, pAGG, pATA, pATT, and pATC) were confirmed by Sanger sequencing.

### 5’ end deletion of the truncated OpIE2 (tIE2) promoter sequence

The tIE2 promoter sequence was split into two fragments tIE2.1 and tIE2.2. The tIE2.1 promoter fragment was obtained by PCR using the OpIE2/F2 forward primer and the reverse primer OpIE2/R2: *5’-TA<u>GAATTC</u>GGCTGCGGTTAGCAACAC-3’*. While the tIE2.2 promoter fragment was obtained by PCR using the forward primer OpIE2/F3: *5’-TAT<u>CTCGAG</u>TACACTACCACACATTGAA-3’* and OpIE2/R1 reverse primer using the same cycling conditions mentioned in the previous section. The tIE2.1 and tIE2.2 promoter fragments were then cloned into pEGFP-N1 vector (Clontech, USA) to produce pCMV/tIE2.1 and pCMV/tIE2.2 vectors, respectively. Also, nine shorter lengths of the truncated OpIE2 (tIE2) promoter were obtained using the same reverse primer (OpIE2/R1) and nine different forward primers listed in Table S2, with the longest fragment of 452 nucleotides from the transcription start site and the shortest of 45 nucleotides from the transcription start site. These fragments were then cloned into pEGFP-N1 vector (Clontech, USA) between *Xho*I and *Eco*RI sites to produce the following recombinant vectors; (pCMV/tIE2-452, pCMV/tIE2-406, pCMV/tIE2-358, pCMV/tIE2-313, pCMV/tIE2-261, pCMV/tIE2-209, pCMV/tIE2-149, pCMV/tIE2-110, and pCMV/tIE2-45).

### Introducing a spacer sequence between CMV and OpIE2 promoters

A 329 bp DNA fragment was isolated from ORF 128 (1044-1372 bp) in the genome of Autographa californica multiple nucleopolyhedrovirus (AcMNPV) insect virus, using the forward primer: *5’-TAT<u>CTCGAG</u>CAGAAAAAAGACGCGCAGG-3*’ and the reverse primer: 5’-*TA<u>GAATTC</u>CCTATTTTCAAATTGTTGC*-3’ with the same protocol and cycling conditions mentioned above. This sequence contained no ATG codons and was cloned between the CMV promoter sequence and tIE2.1 and tIE2.2 fragments in pCMV/tIE2.1 and pCMV/tIE2.2 vectors, respectively. pCMV/tIE2.1 and pCMV/tIE2.2 vectors were cut by NheI restriction enzyme (New England Biolabs-NEB, USA) and subjected to gap-filling by *Phusion* high fidelity Taq polymerase (New England Biolabs-NEB, USA) producing blunt ends. The vector and insert were subjected to blunt end ligation by T4 ligase (New England Biolabs-NEB, USA). Correct ORF orientation in pCMV/329/tIE2.1 and pCMV/329/tIE2.2 was confirmed by colony PCR.

### Cell Line Maintenance and Transfection

*Spodoptera frugiperda*-Sf9 insect cell line was maintained in a monolayer on ExCell-420 insect media (Sigma Aldrich, USA) supplemented with 10% Fetal Bovine Serum (FBS) (Gibco, ThermoFisher Scientific, USA) at 27°C. For the transfection process, Sf9 cells were seeded in 24-well plates at a density of 2×10^5^ cells per well. The transfection mixture was prepared by mixing 2 µg plasmid DNA and 4 µl Cellfectin-II reagent (Invitrogen, ThermoFisher Scientific, USA) to a final volume of 200 µl serum-free media. The transfection mixture was incubated on the cells for 4 h before replacement by supplemented ExCell-420 fresh media; cells were left until examination 48 h post-transfection.

HeLa and HEK-293T mammalian cell lines were maintained in RPMI and DMEM-High glucose (Gibco, ThermoFisher Scientific, USA) media, respectively, supplemented with 10% FBS (Gibco, ThermoFisher Scientific, USA) and incubated with 5% CO_2_ at 37°C. For transfection, cells were seeded in 24-well plates at a density of 50,000 cells/well. The transfection was performed by Lipofectamine 3000 transfection reagent (Invitrogen, ThermoFisher Scientific, USA). For each clone, 0.5 µg DNA were added to the cells. The transfection complexes were prepared in Opti-mem reduced serum media (Gibco, ThermoFisher Scientific, USA) and incubated at room temperature for 30 min before adding to cells. The transfected cells were incubated at 37°C with 5% CO_2_ for 48 h before imaging.

### Fluorescent Microscopy

Cells expressing EGFP protein were visualized 48 h post-transfection by fluorescent microscopy using Leica DMi8 inverted microscope (Leica Microsystems, Germany) at magnification 10x, with an excitation wavelength of 488 nm and an emission wavelength of 509 nm. Photos were taken by The Leica Application Suite X (LAS X) software. Images were processed using ImageJ software (Rueden 2017) to estimate the fluorescnce intensities of cells by calculating the mean gray value after background subtraction.

### Total RNA extraction

Total RNA was extracted 48 h post-transfection using the RNeasy Mini Kit (Qiagen) following the manufacturer’s instructions. From a 24-well plate, a total of 2×10^5^ cells/well were collected to extract the total RNA. The extracted RNA concentration and purity were determined by NanoDrop (ThermoFisher Scientific, USA). During extraction, the RNA was treated on-column by DNase I RNase-free enzyme (Qiagen) according to the kit’s protocol.

### cDNA synthesis and Quantitative Real-time PCR

The cDNA was synthesized by random primers using High Capacity cDNA Reverse Transcription Kit (ThermoFisher Scientific). Two µg of the extracted RNA were used per reaction. Quantitative real-time PCR was performed on Quantstudio 12K Flex (ThermoFisher Scientific, USA) using PowerUP SYBR green master mix (ThermoFisher Scientific, USA) and the following conditions: 50°C for 2 min, 95°C for 2 min then 40 cycles (95°C for 1 s and 60°C for 30 s). The primers used to amplify the EGFP gene were as follows: *EGFP/F: 5’-CTGCTGCCCGACAACCA-3’* and *EGFP/R: 5’-TGTGATCGCGCTTCTCGTT-3’*. β-Actin gene was used as an internal control using *q β-Actin-F: 5’-CATGGAGTCCTGTGGCATC-3’ and q β-Actin-R: 5’-CAGGGCAGTGATCTCCTTCT-3’* primers.

### In-gel fluorescence

Total protein was extracted from the cells 48 h post-transfection by complete lysis-M buffer (Merck, USA). Twenty-five mg of total protein were loaded per well in a 12% SDS PAGE. EGFP fluorescence was visualized using ChemiDoc-XRS+ and Image Lab software (Bio-Rad, USA) at excitation/ emission: 493/518 nm.

### Fluorescent-Activated Cell Sorter (FACS)

The population of fluorescent cells expressing EGFP was determined by FACS. Cells were collected 48 h post-transfection and resuspended in PBS buffer. Flow cytometry was carried out using FACSCalibur (Becton Dickinson, USA) following standard procedures with CellQuest Pro Software (Becton Dickinson, USA). Data analysis was performed using FlowJo v. 10.2 software (Treestar, USA). For each sample, 10,000 cells were analyzed. The mean fluorescence intensity (MFI) was calculated using normalized X Geo Mean values.

## RESULTS

### Construction of the mammalian/insect shuttle vectors

The shuttle vectors were designed based on the pEGFP-N1 vector backbone, in which different versions of the OpIE2 promoter (truncated and ATG mutated) were cloned downstream of the CMV promoter (Figure 1A). The wild-type OpIE2 promoter sequence contains seven ATG triplets that may act as potential start codons. Six of them are located in the first 70 bp. A truncated version of the OpIE2 promoter (tIE2) was created, that lacks the first six ATGs leaving only one ATG at −75 position from the transcription start site (Figure 1B). The wild-type OpIE2 promoter (561 bp) and truncated OpIE2 (tIE2) promoter (491 bp) were each cloned separately into the pEGFP-N1 vector, downstream from the CMV promoter and upstream from the EGFP gene to produce pCMV/IE2 and pCMV/tIE2, respectively (Figure 2). A control vector is also constructed in which the CMV promoter was removed from pCMV/tIE2 to produce pCMV/tIE2ΔCMV. This construct was later used as a positive control for EGFP gene expression in Sf9 cells.

**Figure 1.**
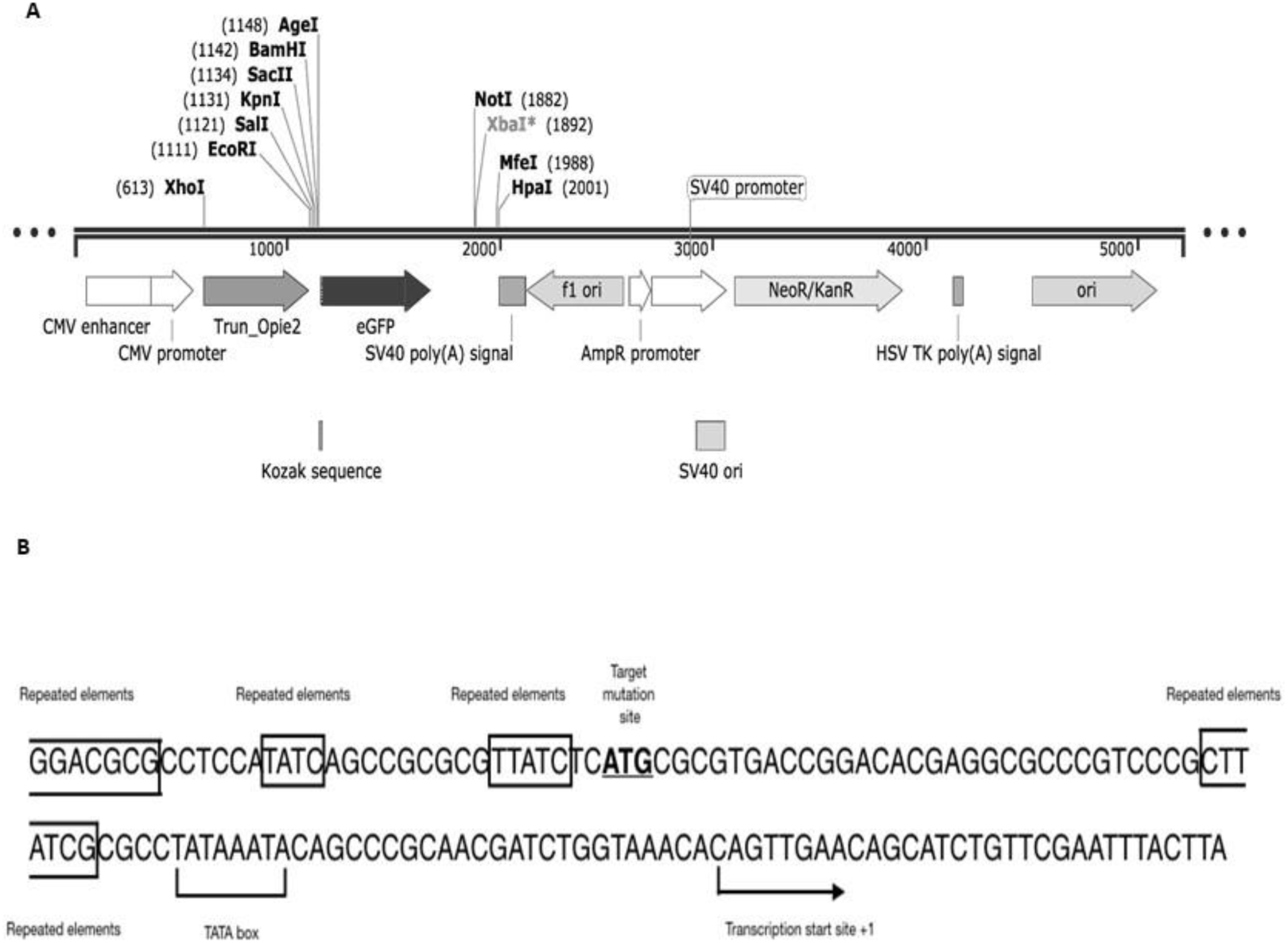
A schematic diagram of the shuttle vectors construction. (A) The constructed shuttle vector is based on the pEGFP-N1 vector backbone. It has the CMV constitutive promoter that can induce EGFP expression in mammalian cells. The truncated viral promoter (tIE2) was cloned downstream of the CMV promoter to direct EGFP expression in insect cells. (B) The last 140 bases at the 3’ end of the tIE2 promoter, showing ATG triplet (underlined) at position −75 of the OpIE2 promoter that could be a potential translation start codon. Some repeated element of the promoter in these 140 bases are marked in triangles. The TATA box and transcription start site are marked as well.

**Figure 2.**
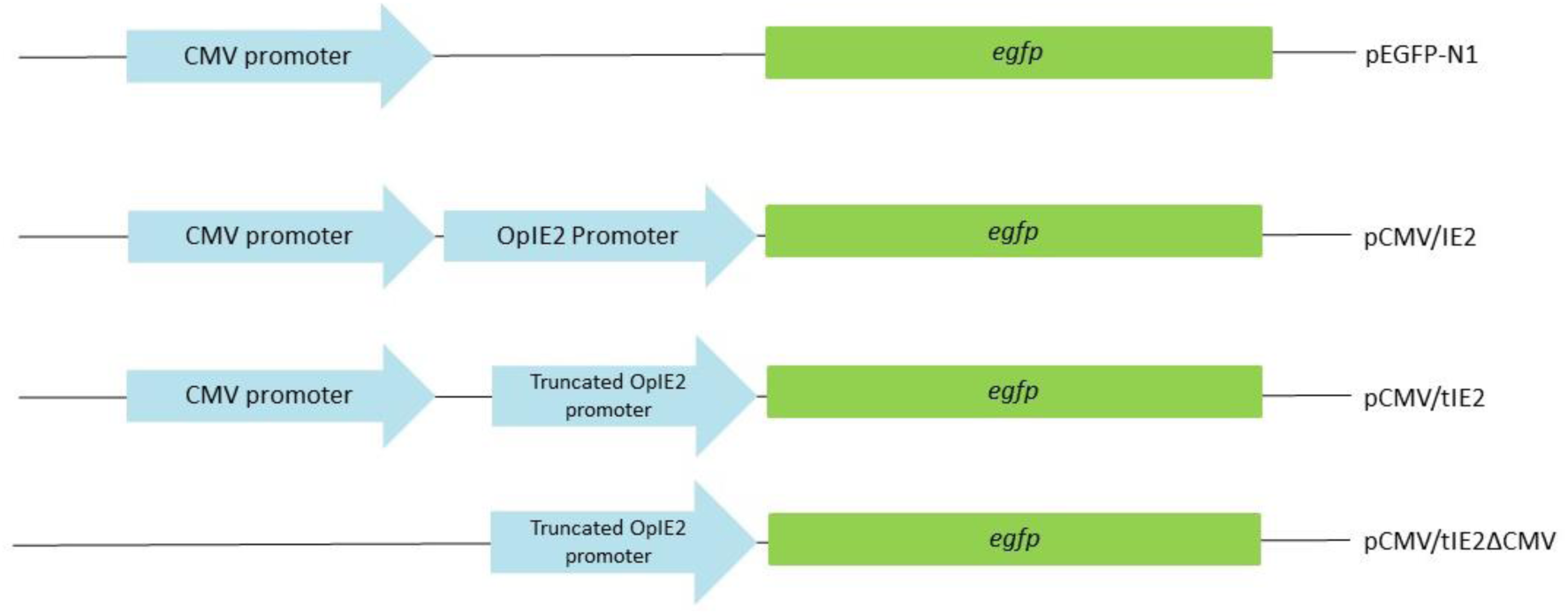
Schematic diagram of the shuttle and control vectors. The figure depicts the arrangement of the CMV promoter and different versions of the OpIE2 promoter followed by the EGFP reporter gene. Seventy nucleotides from the 5’ end of the OpIE2 promoter were removed in the truncated promoter (tIE2). pEGFP-N1 is the vector used as a backbone for the other vectors. pCMV/IE2 and pCMV/tIE2, are pEGFP-N1 with the whole IE2 or the truncated IE2, respectively. A control clone was constructed by removing the CMV promoter from pCMV/tIE2 (pCMV/tIE2ΔCMV).

### Prediction of transcription-factors’ binding sites in tIE2 promoter sequence

The truncated OpIE2 (tIE2) promoter lacks six of the seven ATGs present in the wild type OpIE2 promoter. Therefore, mutations should be conducted to the only ATG left in the tIE2 sequence at −75 position to produce a tIE2 promoter without any ATG in its sequence. However, before conducting the site-directed mutagenesis, computational validation was performed to predict that mutations at the targeted ATG trinucleotide would not alter the promoter’s activity. Due to the lack of information about the transcription factors that may bind to the tIE2 promoter, a prediction model for the full pattern of transcription factors’ binding sites (TFBSs) was established *in silico* using the Match analysis tool (Kel 2003) in the TRANSFAC (Matys 2003) manually curated database. The resulting transcription factor binding pattern, based on insects’ profile, was used to build positional weight matrices (PWMs) that were used to search for possible TFBSs. Most of the transcription factors found were related to *Drosophila melanogaster*. As shown in Figure 3, 16 different transcription factors were detected to bind to the truncated OpIE2 (tIE2) promoter sequence; some of them bind to repeated sequences that are scattered throughout the promoter sequence (Table S3).

**Figure 3.**
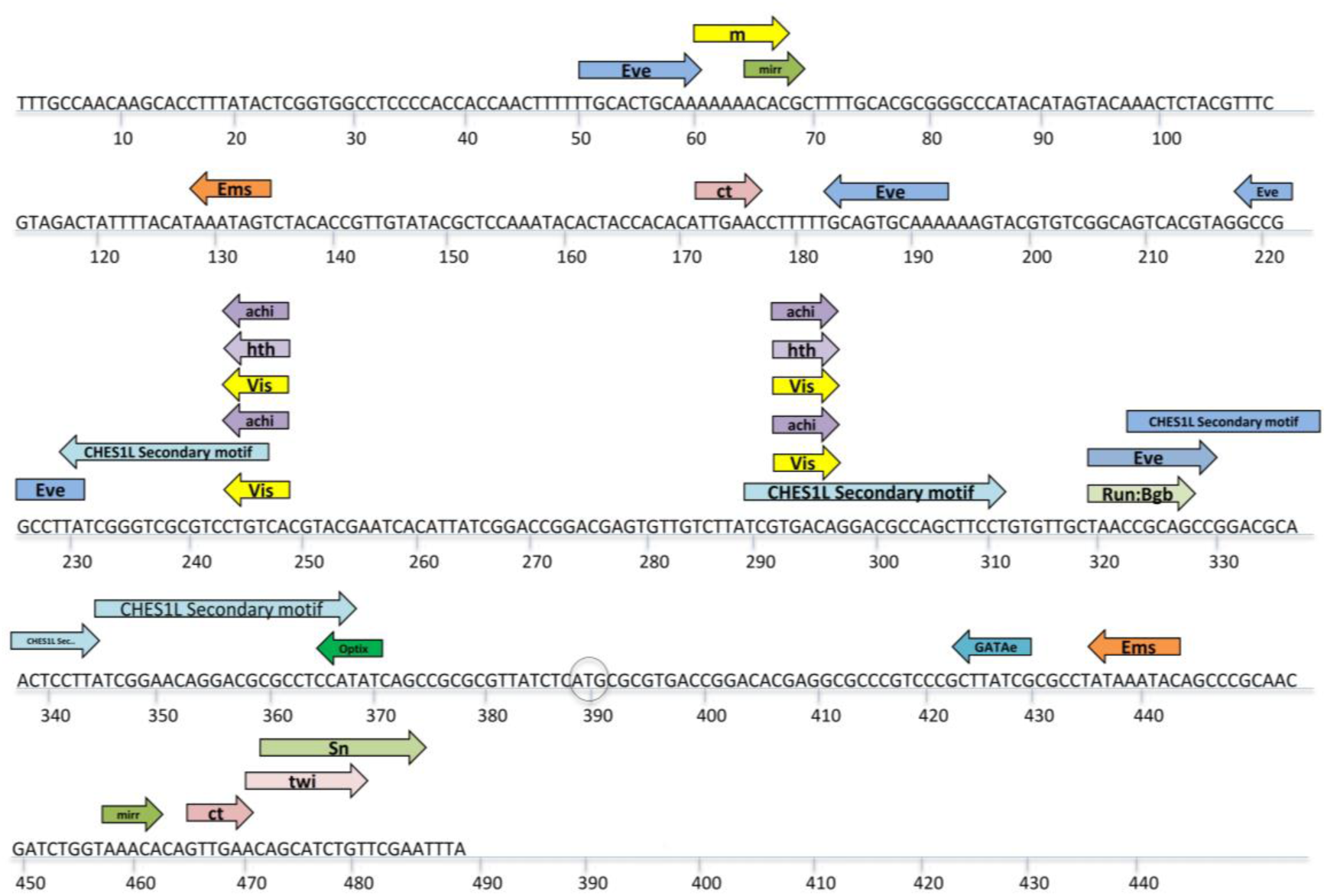
*In silico* analysis of tIE2 promoter sequence. The Transcription factor binding pattern of tIE2 promoter sequence and the putative TFBS on the truncated version of the wild type OpIE2 (tIE2) promoter sequence. The model is built using TRANSFAC database Match tool that utilizes insects’ profile to build PWMs from the model. The target site for mutation (bases between the region: 380 to 390 bp, pointed by a black circle) showed no TF binding. The ATG at −75 bp from the transcription start site is marked with a black circle.

The ATG at −75 position was not shown to be a target for any TFBSs even when all nine possible nucleotide substitutions to alter the ATG codon (<u>T</u>TG, <u>C</u>TG, <u>G</u>TG, A<u>A</u>G, A<u>C</u>G, A<u>G</u>G, AT<u>A</u>, AT<u>C</u>, and AT<u>T</u>) were used in the analysis.

### *In silico* analysis of the chimeric CMV/tIE2 promoter sequence showed no change in the transcription factors binding to the tIE2 sequence

The CMV promoter has minimal activity in uninfected insect cell lines (Cheng 2004). To ensure that the presence of the CMV promoter upstream the tIE2 promoter will not interfere with the recruitment of the necessary transcription factors on the tIE2 promoter, another binding pattern was created using the chimeric CMV/tIE2 promoter sequence. The fused promoters’ sequences were scanned using the same criteria on the Match analysis tool in insect profile (Figure 4). A total of 25 different potential insect transcription factors were recognized to bind to the CMV promoter sequence. At the enhancer region (−500 to −300 from the transcription start site), which is part of the CMV promoter in pEGFP-N1, 13 different transcription factors were predicted; the rest of the transcription factors have their binding sites on the proximal sequence of the CMV promoter. Another 15 transcription factors were associated with their binding sites from −300 to +1. It was noticed that the CMV promoter sequence recruited a completely different set of insect transcription factors when compared to those on the tIE2 promoter sequence. Furthermore, there was no change observed in the binding pattern, score, or directions of the transcription factors on the tIE2 promoter after including the CMV promoter sequence.

**Figure 4.**
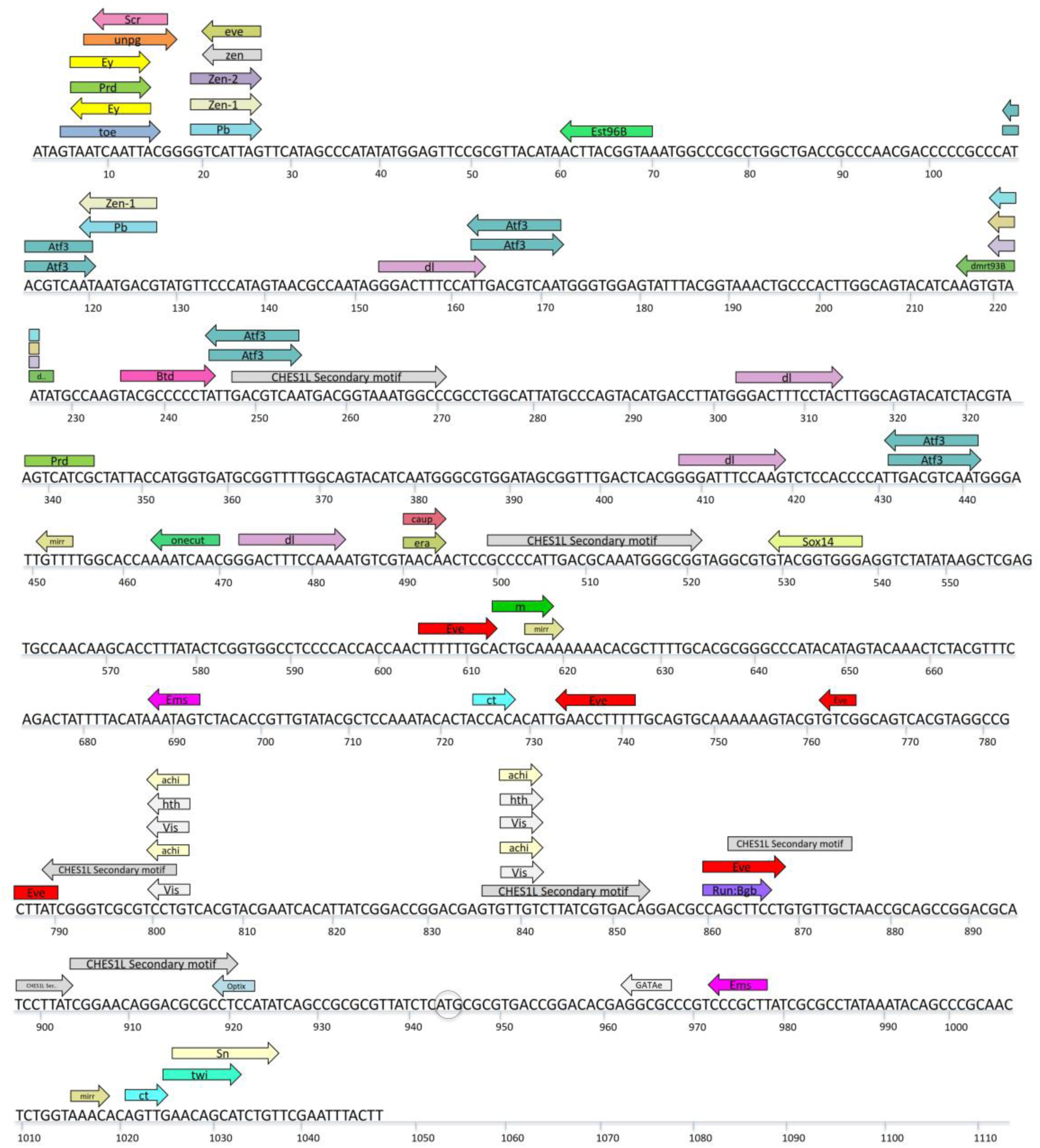
*In silico* analysis of the chimeric CMV/tIE2 promoter. Match tool analysis results for the chimeric CMV/tIE2 promoter sequence. No change was observed in the binding pattern, score, or directions of the transcription factors on the tIE2 promoter sequence when combined with the CMV promoter sequence. The ATG at −75 bp from the transcription start site is marked with black circle.

### The chimeric CMV/tIE2 promoter worked efficiently in Sf9 cells but not in HEK293T cells

pCMV/tIE2 was introduced to Sf9 and HEK-293T cells, and cells were visualized 48 h post-transfection under fluorescence microscopy (Figure 5). tIE2 activity (indicated by the fluorescence intensity) in Sf9 cells was not compromised after removing the first 70 bp from the wild-type OpIE2 promoter sequence (Figure 5A and B). On the other hand, both full-length and truncated OpIE2 (tIE2) sequences seemed to affect the upstream promoter (CMV) negatively in HEK-293T cells in comparison to pEGFP-N1 (Figure 5C and D). However, there was a slight increase in EGFP fluorescence intensity upon transfection with pCMV/tIE2 compared to pCMV/IE2, which may have been due to the lack of the six ATGs in the tIE2 promoter sequence. Therefore, we started testing the effect of the mutation in the only remaining ATG triplets in the tIE2 promoter sequence at −75 position.

**Figure 5.**
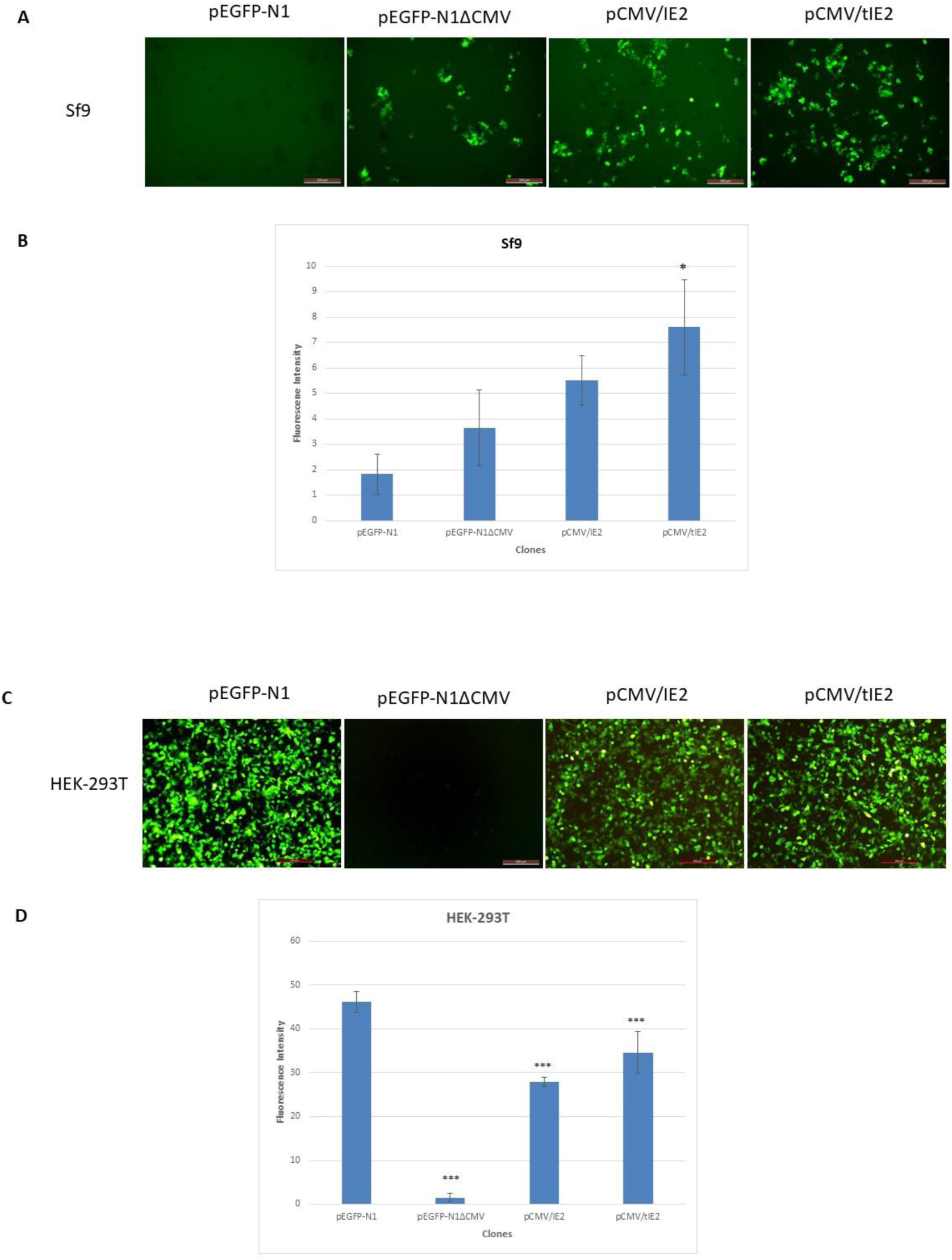
Truncated OpIE2 promoter activity in Sf9 and HEK-293T cells. (A) EGFP fluorescence in Sf9 cells transfected with pCMV/IE2, pCMV/tIE2 vectors in addition to pEGFP-N1 and pEGFP-N1ΔCMV vectors as controls. (B) A graph representing the numerical values of the Sf9 cell images in Figure 5A. (C) EGFP fluorescence in HEK-293T cells transfected with pCMV/IE2, pCMV/tIE2 vectors in addition to pEGFP-N1 and pEGFP-N1ΔCMV vectors as controls. (D) A graph representing the numerical values of the HEK-293T cell images in Figure 5C. The structure of the vectors is represented in Figure 2. Cells were harvested 48 h post-transfection and imaging was done by inverted fluorescent microscopy. The numerical values were obtained using ImageJ software. ** represents significance of P<0.01, and *** represents significance of P< 0.001.

### The SDM versions of the CMV/tIE2 promoter showed similar activity to CMV/tIE2 promoter in both insect and mammalian cells

Site-directed mutations were conducted to alter the remaining ATG triplet at −75 position in the tIE2 sequence. All possible mutations were adopted (AAG, ACG, AGG, ATA, ATT, ATC, TTG, CTG, and GTG) as shown in Figure 6A. These mutations were made to the pCMV/tIE2 vector to produce nine different vectors (pAAG, pACG, pAGG, pATA, pATT, pATC, pTTG, pCTG, and pGTG). All nine vectors were individually transfected into Sf9 and HEK-293T cells to test their EGFP expression levels. For in-gel fluorescence analysis, total protein was extracted and run on SDS-PAGE, then imaged to detect the green fluorescence intensity. All nine mutant versions of pCMV/tIE2 were capable of expressing EGFP efficiently in Sf9 cells (Figure 6B), but not in HEK-293T cells, similar to the original pCMV/tIE2 (Figure 6C). Some in-gel fluorescent bands were more intense than bands in the original pCMV/tIE2 in figures 6 B and C. However, this intensity was observed in some but not all mutations, which indicated that it is not related to the absence of ATG triplets specifically.

**Figure 6.**
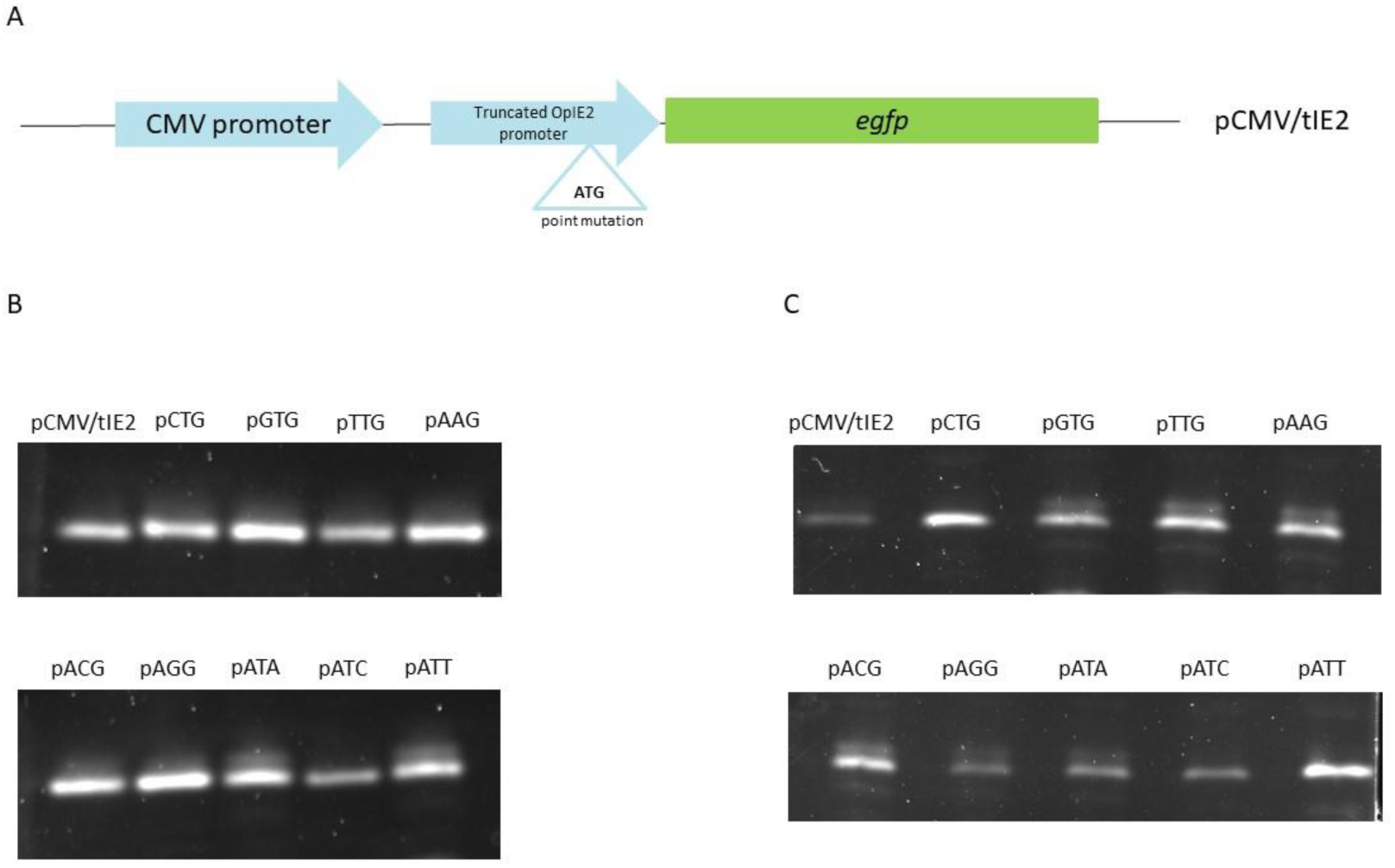
The effect of SDM created in CMV/tIE2 chimeric promoter on gene expression. (A) The position of SDM created in the truncated OpIE2 promoter at position −75. The nine mutated constructs were referred to as (pCTG, pGTG, pTTG, pAAG, pACG, pAGG, pATA, pATC, and pATT). (B) In-gel fluorescence images of the nine mutated vectors expression patterns in Sf9 cells. (C) In-gel fluorescence images of the nine mutated vectors in HEK-293T cells. Protein was extracted from transfected cells 48 h post-transfection and run on 12% SDS PAGE.

### The reduction in the activity of the chimeric CMV/tIE2 promoter in mammalian cells was linked to transcription, not translation

Since the mutated pCMV/tIE2 vector (with no ATG triplets in tIE2 promoter sequence) did not restore the original CMV promoter activity in mammalian cells, the EGFP transcript level was measured by RT-qPCR for pCMV/tIE2 and pEGFP-N1 vectors in HEK-293T cells. The EGFP transcript level was reduced by by a factor of five in pCMV/tIE2 in comparison to pEGFP-N1 (Figure 7).

**Figure 7.**
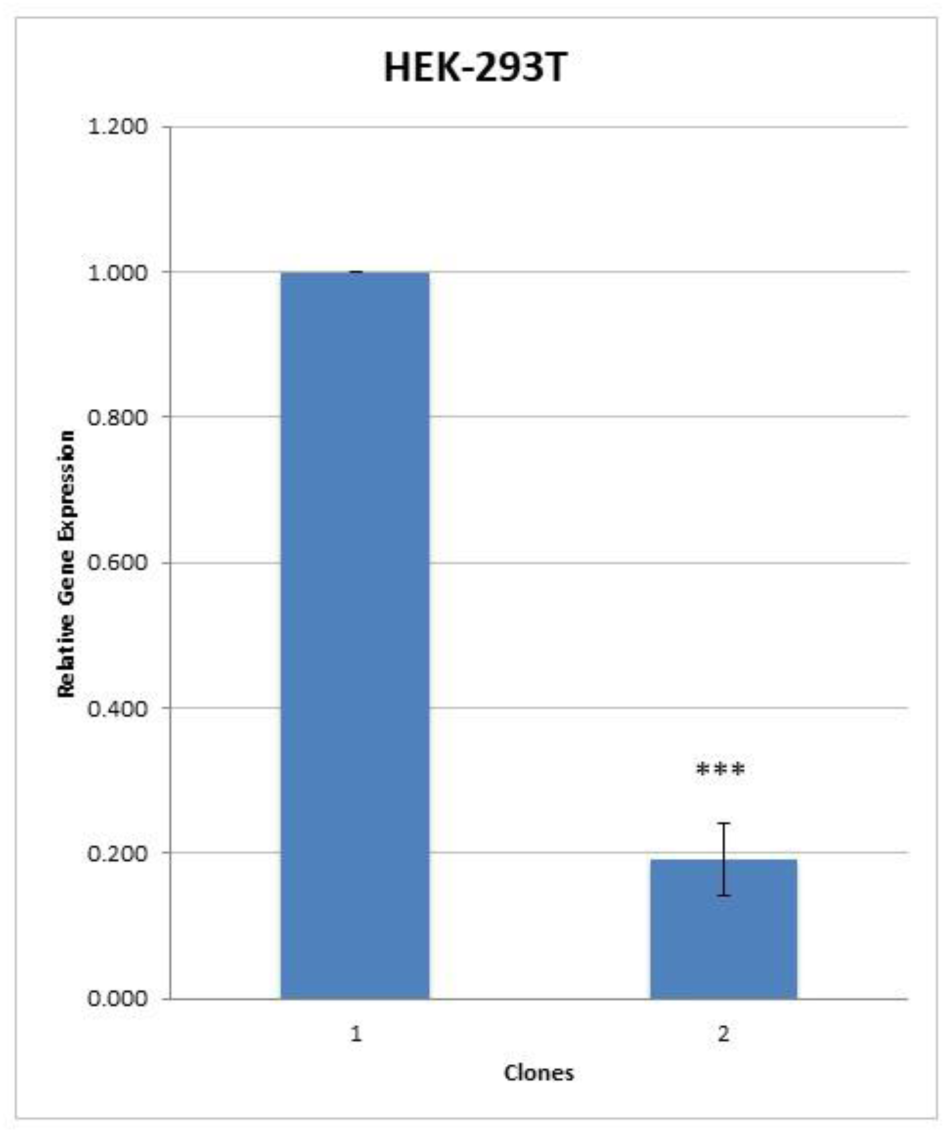
Quantification of pCMV/tIE2 expression in HEK-293T cells. EGFP was expressed by pEGFP-N1 and pCMV/tIE2 vectors in HEK-293T cells. The level of gene expression was measured by RT-qPCR 48 h post-transfection. Each data value was obtained from the mean of three independent replicates. *** represents statistical significance of P< 0.001.

### The tIE2 promoter sequence seems to have a repressive effect on CMV promoter activity

The tIE2 promoter was divided into two fragments of 329 bp each, with an overlapping region of 170 bp. The two fragments tIE2.1 and tIE2.2 were cloned onto pEGFP-N1 to produce pCMV/tIE2.1 and pCMV/tIE2.2 vectors, respectively (Figure 8A). A control vector (pCMV/329) was constructed by cloning 329 bp of ORF 128 of Autographa californica nucleopolyhedrovirus (AcMNPV) in pEGFP-N1 vector downstream CMV promoter. These 329 bp contain no ATG triplets and do not contain any regulatory sequences (Figure 8A). pCMV/tIE2.1 and pCMV/tIE2.2 were transfected into HEK-293T cells to assess the repressive effect of each fragment on the CMV promoter. In Sf9 cells, pCMV/tIE2.2 maintained optimum activity, while pCMV/tIE2.1 showed low EGFP fluorescence intensities (Figure 8B, C, and H). In HEK-293T cells, pCMV/tIE2.1 did not show notable enhancement in CMV promoter activity compared to pCMV/tIE2, while pCMV/tIE2.2 showed higher EGFP fluorescence intensities (Figure 8D, E, and H). For further confirmation and generalization of this result, pCMV/tIE2.1 and pCMV/tIE2.2 were tested in HeLa cells as well (Figure 8F, G, and H).

**Figure 8.**
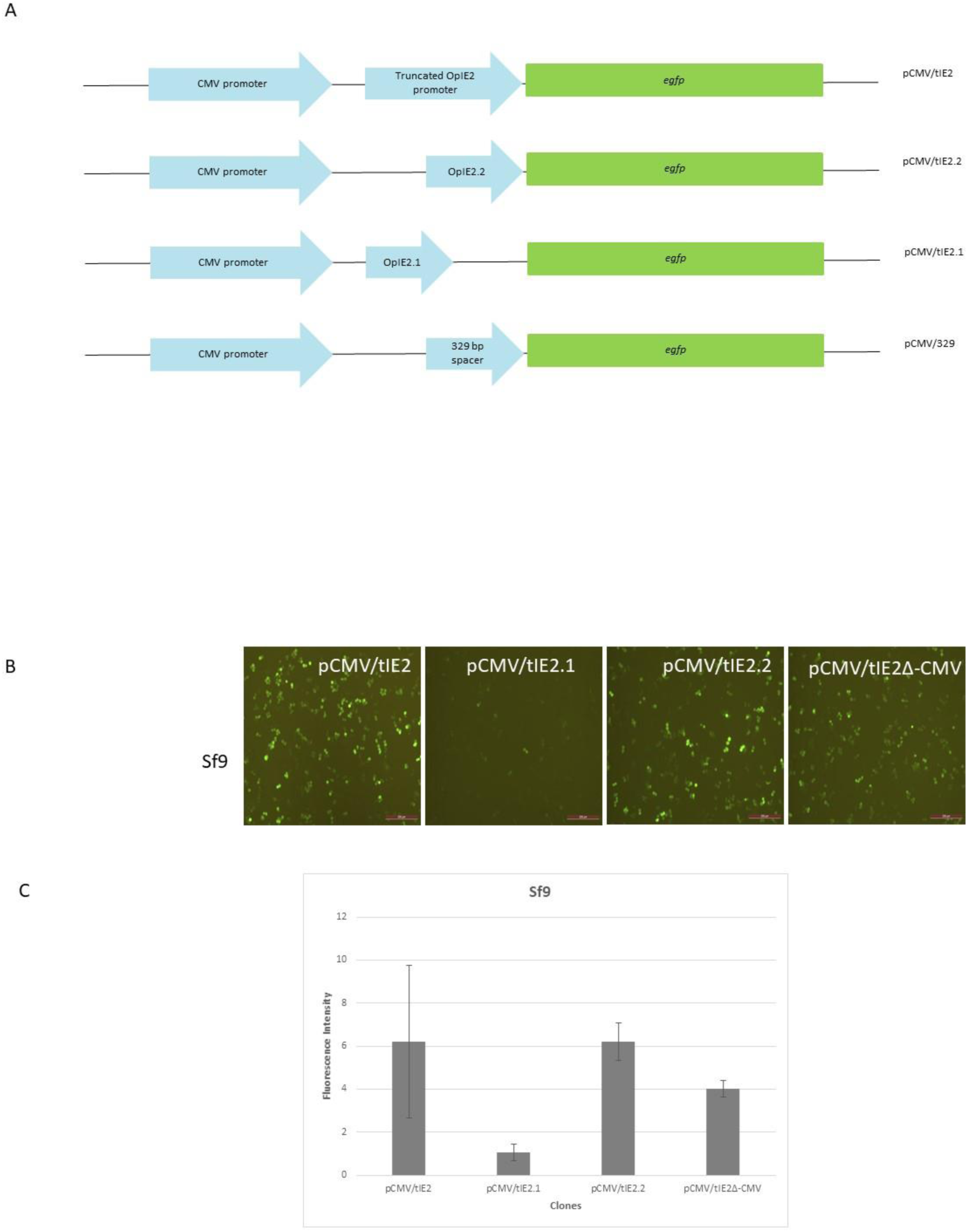
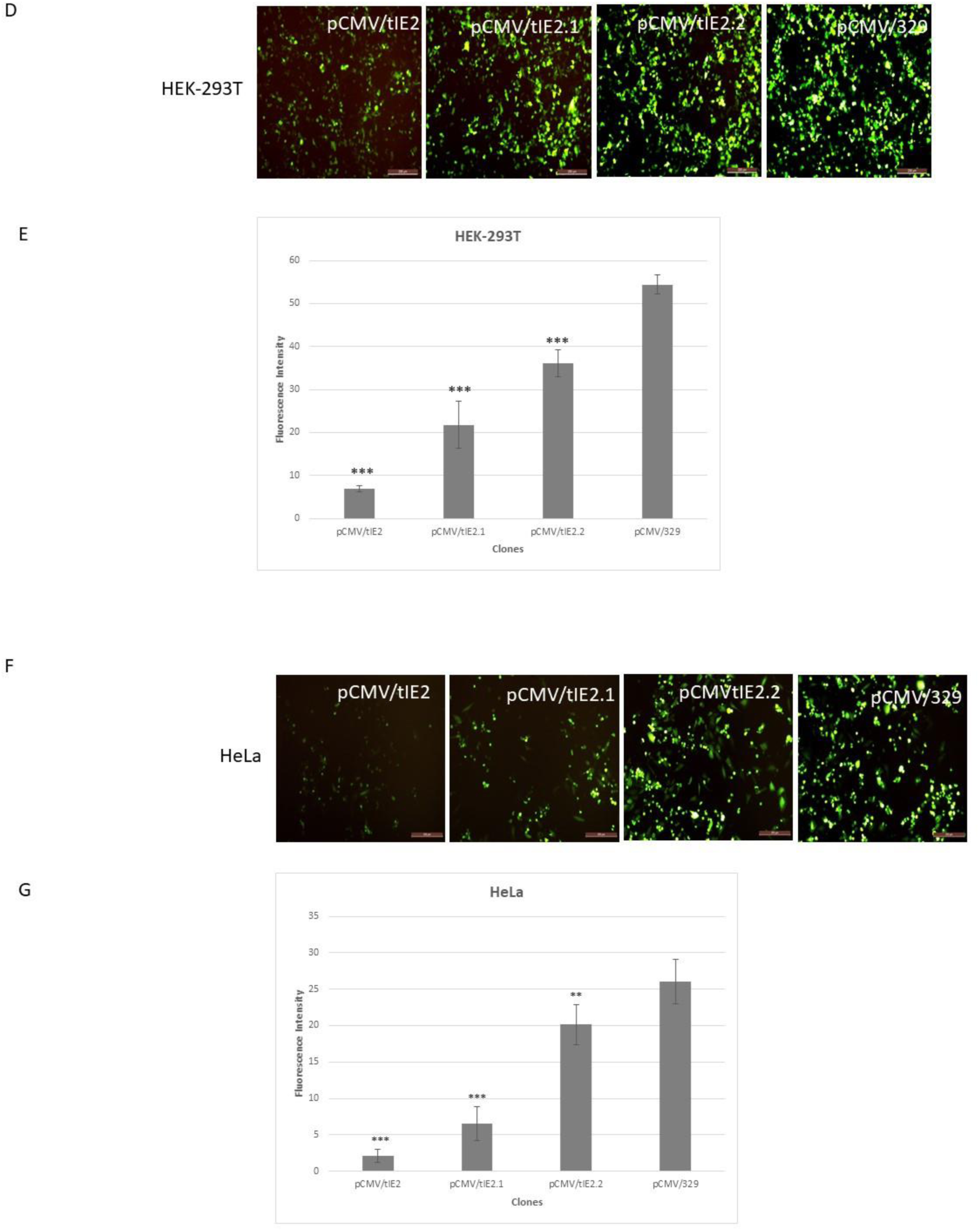
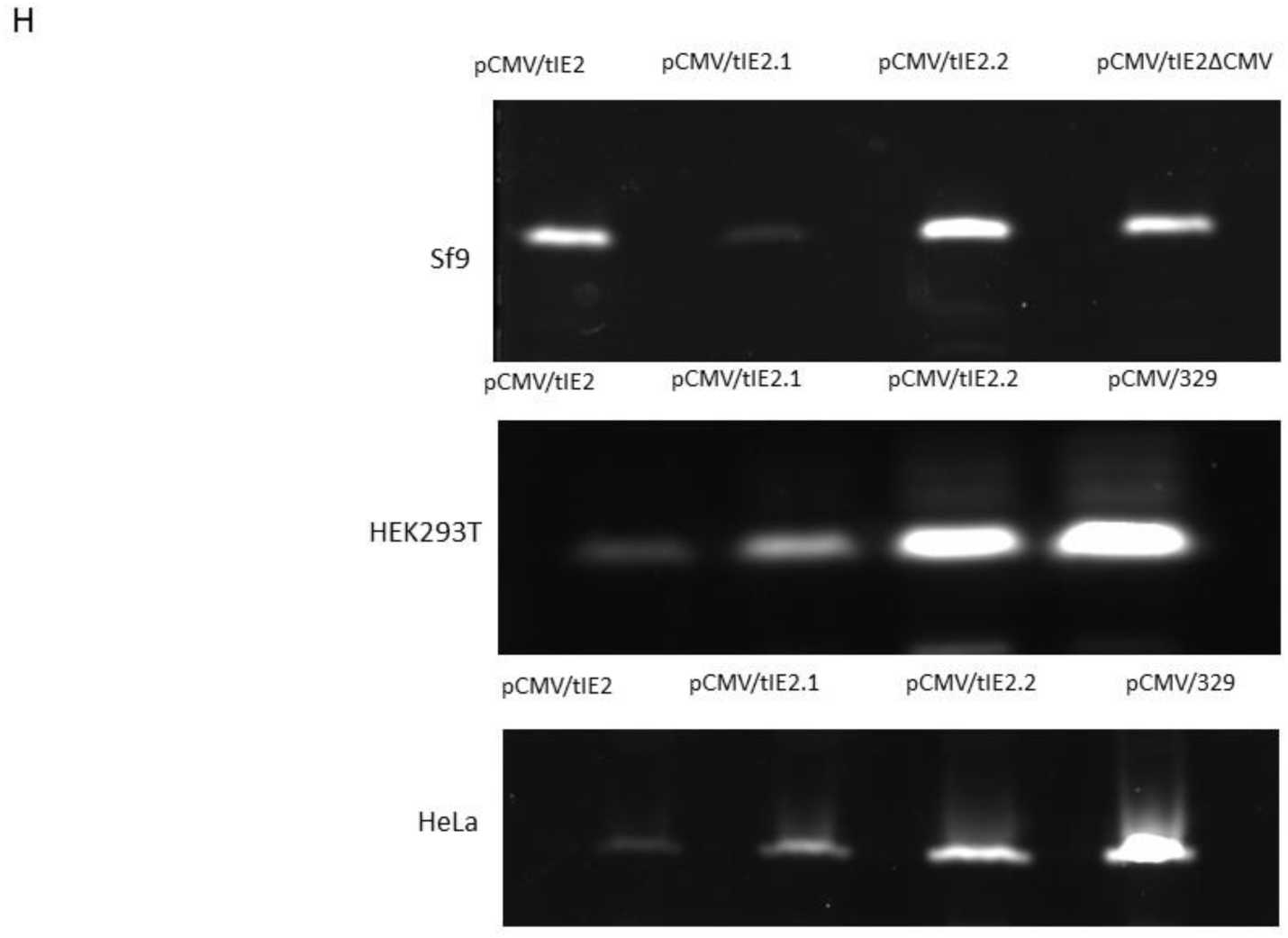
Comparison of EGFP expression driven by the 5’ portion or the 3’ portion of the tIE2 promoter in Sf9, HEK-293T, and HeLa cells. (A) pCMV/tIE2 (truncated OpIE2 (tIE2), 70 bp were removed from the OpIE2 5’ end), pCMV/tIE2.1 (329 bp at the 5’ end of the tIE2 promoter), pCMV/tIE2.2 (329 bp at the 3’ end of the tIE2 promoter), and pCMV/329 (a 329 bp baculovirus ORF sequence with no ATG codons). (B) Expression of pCMV/tIE2, pCMV/tIE2.1, pCMV/tIE2.2, pCMV/tIE2ΔCMV in Sf9 cells. (C) A graph representing fluorescence intensity values for Sf9 cells in Figure 8B. (D) EGFP expression by pCMV/tIE2, pCMV/tIE2.1, pCMV/tIE2.2, pCMV/329 in HEK-293T cells. (E) A graph representing fluorescence intensity values for HEK-293T cells in Figure 8D. (F) EGFP expression by pCMV/tIE2, pCMV/tIE2.1, pCMV/tIE2.2, pCMV/329 in HeLa cells. (G) A graph representing fluorescence intensity values for HeLa cells in Figure 8F. All numerical values were obtained using ImageJ software. ** represents significance of P<0.01, and *** represents significance of P< 0.001. (H) Proteins were extracted from transfected cells 48 h post-transfection and run on 12% SDS PAGE for in-gel fluorescence analysis.

### Separating the two promoters did not have a significant effect on the upstream promoter activity in mammalian cells

The 329 bp ORF was cloned upstream the OpIE2.1 and OpIE2.2 promoters and downstream the CMV promoter in both pCMV/tIE2.1 and pCMV/tIE2.2 vectors, respectively (Figure 9A) in an attempt to enhance CMV promoter activity by separating it from the tIE2 promoter sequence. A modest enhancement in the EGFP fluorescence intensity was noticed after pCMV/329/tIE2.1 and pCMV/329/tIE2.2 transfection in HEK-293T cells (Figure 9B and C).

**Figure 9.**
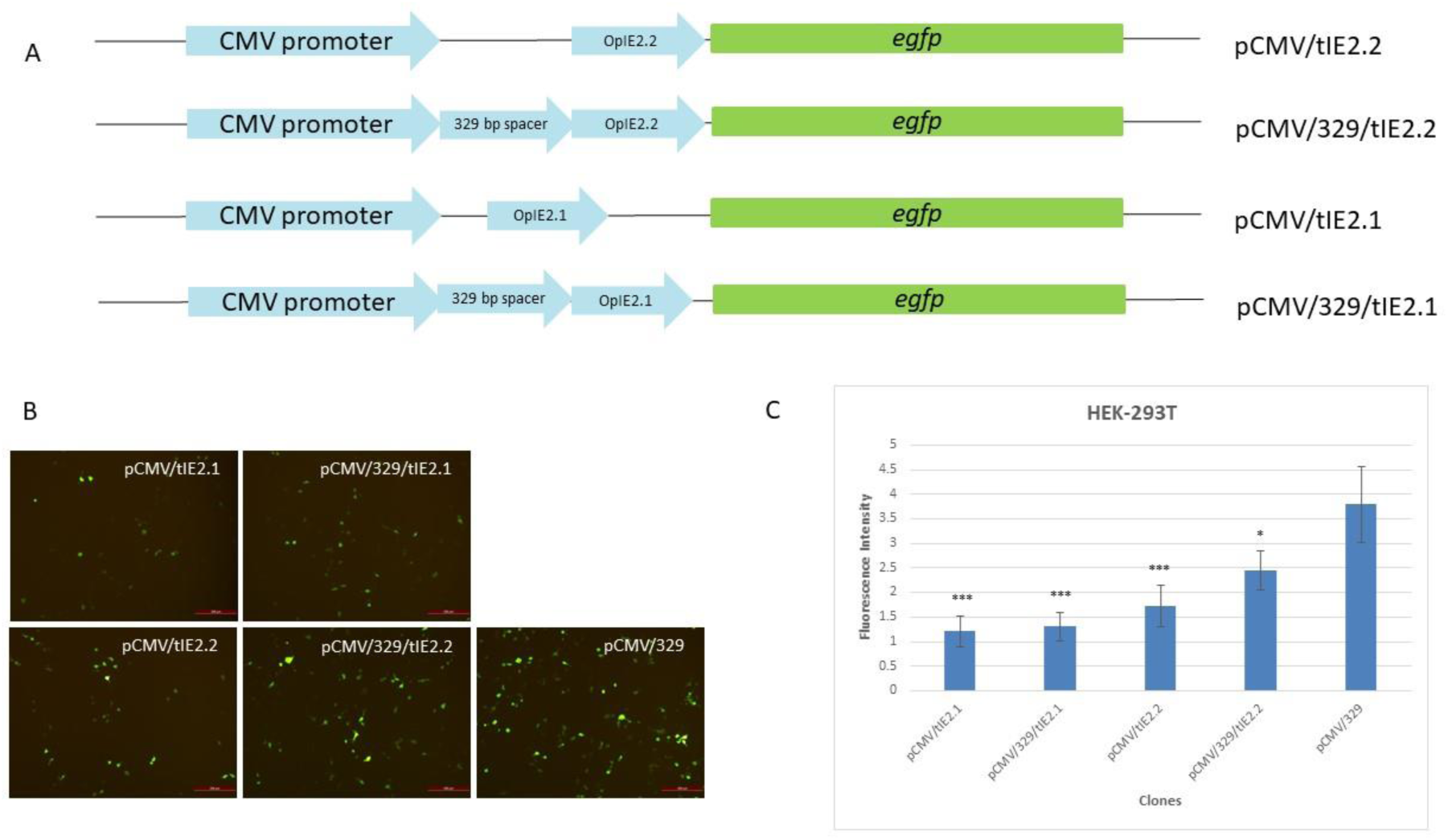
Comparison of EGFP expression driven by the 5’ portion or the 3’ portion of the tIE2 promoter with and without a spacer in HEK-293T cells. (A) A 329 bp ORF with no ATG triplets was cloned upstream of the OpIE2 promoter and downstream the CMV promoter in the pCMV/tIE2.1 and pCMV/tIE2.2 clones to create a space between CMV and OpIE2 promoters. (B) HEK-293T cells were transfected with pCMV/tIE2.1, pCMV/329/tIE2.1, pCMV/tIE2.2, and pCMV/329/tIE2.2 clones, with pCMV/329 as a control. (C) A graph representing fluorescence intensity values for HEK-293T cells in Figure 9B. These values were obtained using ImageJ software. * represents significance of P<0.05, and *** represents significance of P< 0.001.

### 5’ end deletion analysis of the tIE2 promoter revealed inhibitory sequences compromising CMV promoter activity in HEK-293T and HeLa cell lines

The truncated OpIE2 promoter (tIE2) was further truncated using different forward primers that each eliminated 50 bp from the 5’ end of the tIE2 promoter to produce pCMV/tIE-452, pCMV/tIE-406, pCMV/tIE-358, pCMV/tIE-313, pCMV/tIE-261, pCMV/tIE-209, pCMV/tIE-149, pCMV/tIE-110, pCMV/tIE-45 clones (Figure 10A). After transfection in Sf9 cells, clones pCMV/tIE-452, pCMV/tIE-406, and pCMV/tIE-358 (given a purple color code, Figure 10A) showed high fluorescence intensities, clones pCMV/tIE-313, pCMV/tIE-261, and pCMV/tIE-209 (given a green color code, Figure 10A) showed fair fluorescence intensities, while clones pCMV/tIE-149, pCMV/tIE-110, and pCMV/tIE-45 (given an orange color code, Figure 10A) showed no fluorescence (Figure 10B and C). On the contrary, in HEK-239T and HeLa cells, clones pCMV/tIE-452, pCMV/tIE-406, pCMV/tIE-358 showed low fluorescence levels, clones pCMV/tIE-313, pCMV/tIE-261, pCMV/tIE-209, showed fair fluorescence levels, whereas clones pCMV/tIE-149, pCMV/tIE-110, pCMV/tIE-45 showed high fluorescence levels (Figure 10D-G).

**Figure 10.**
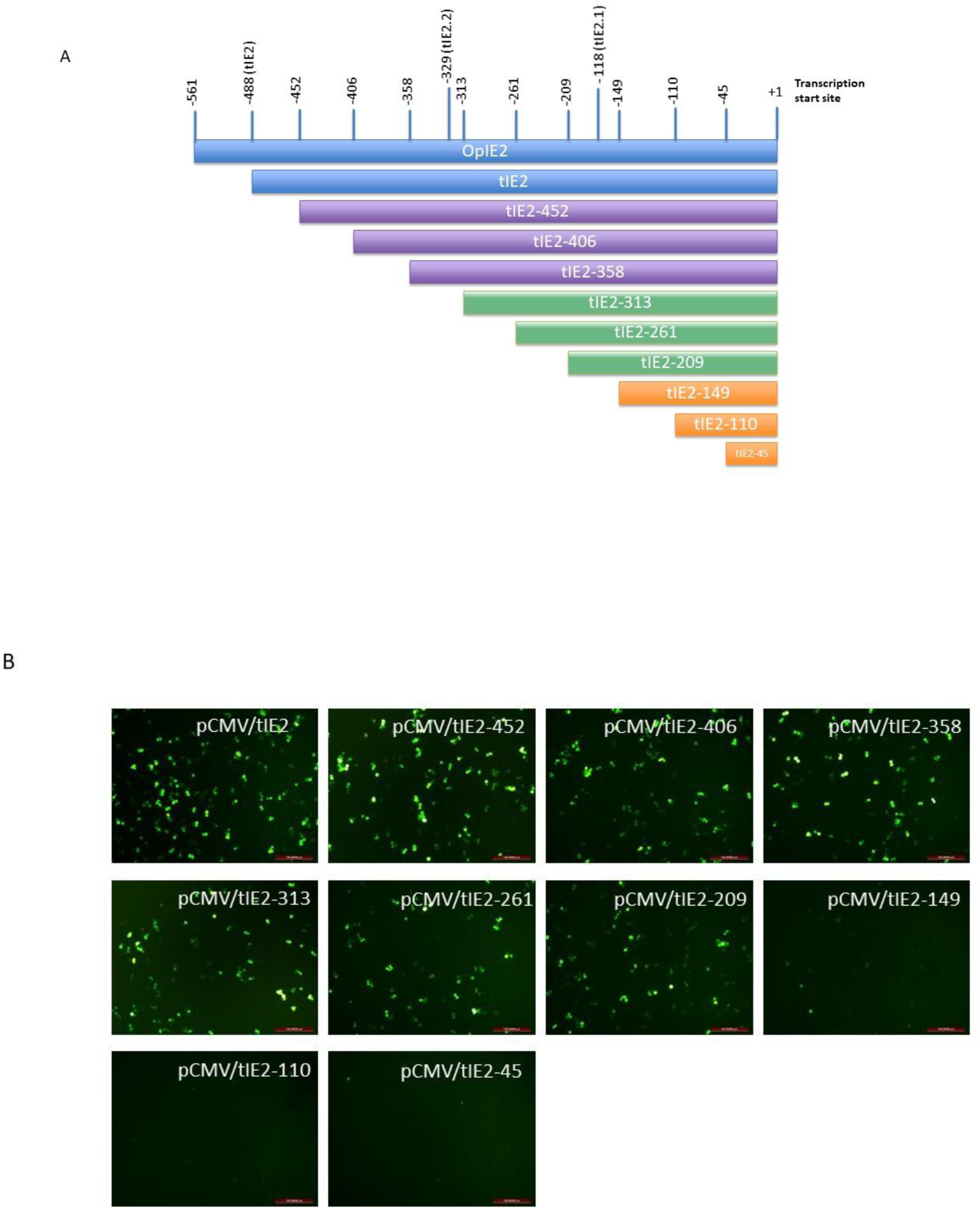
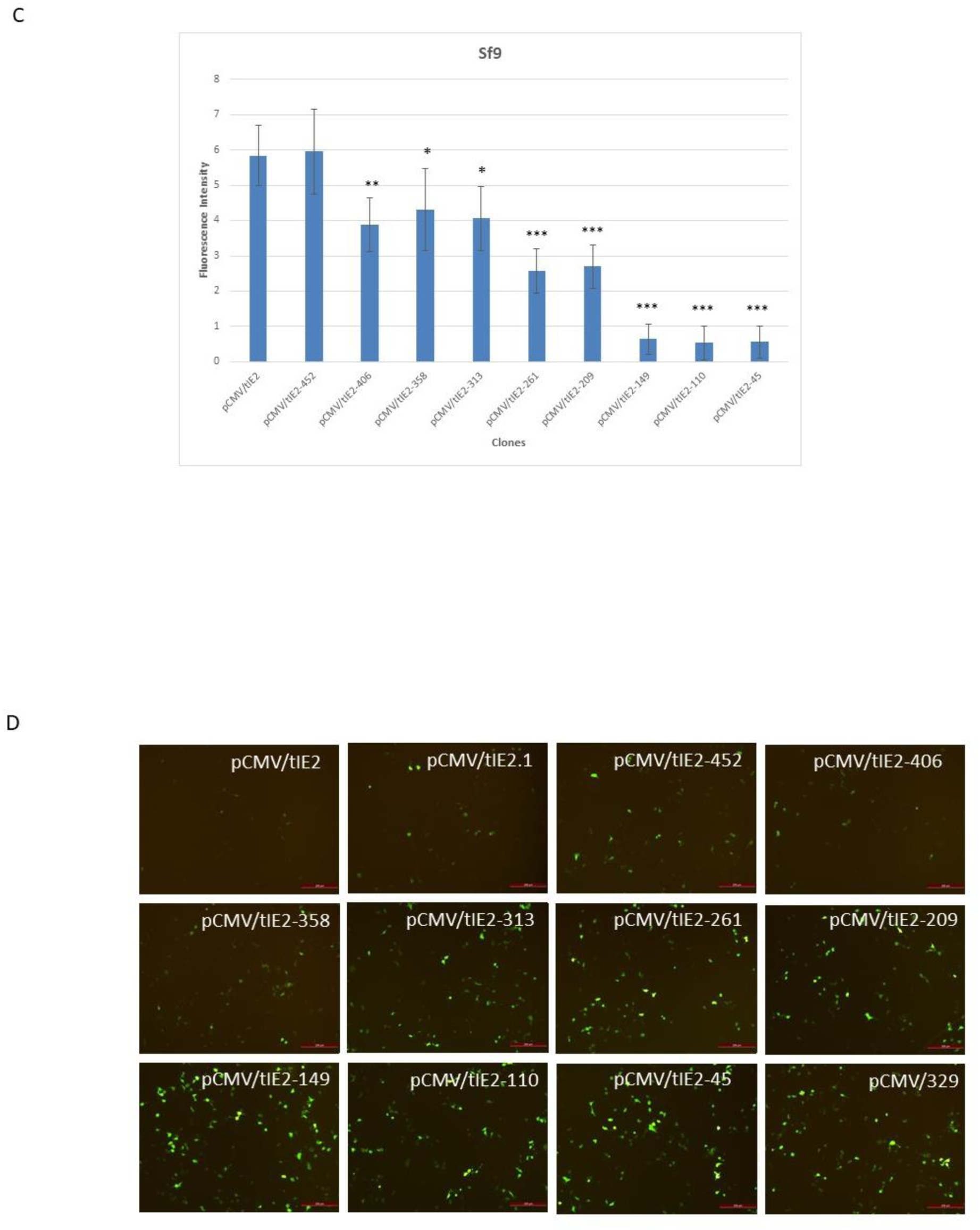
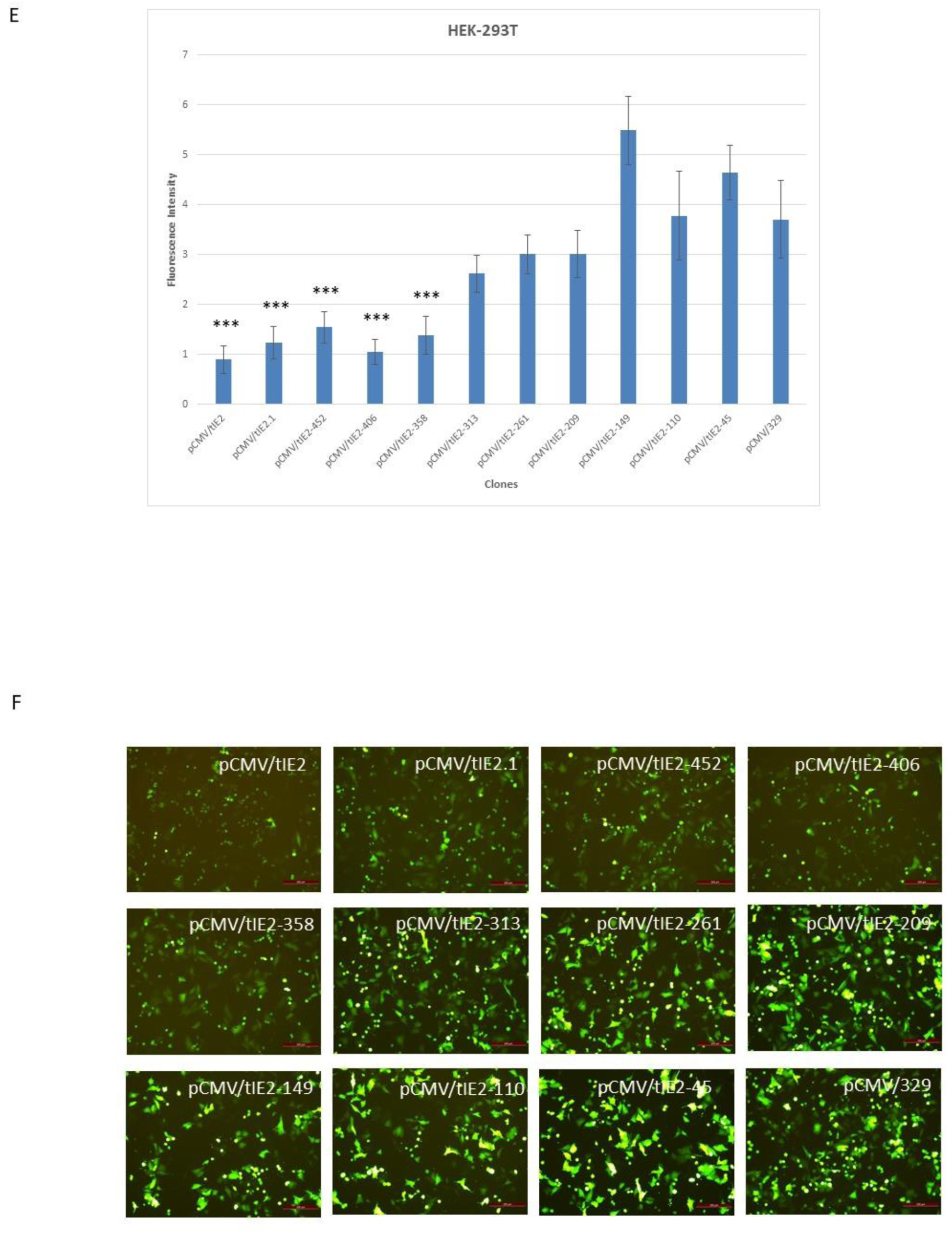
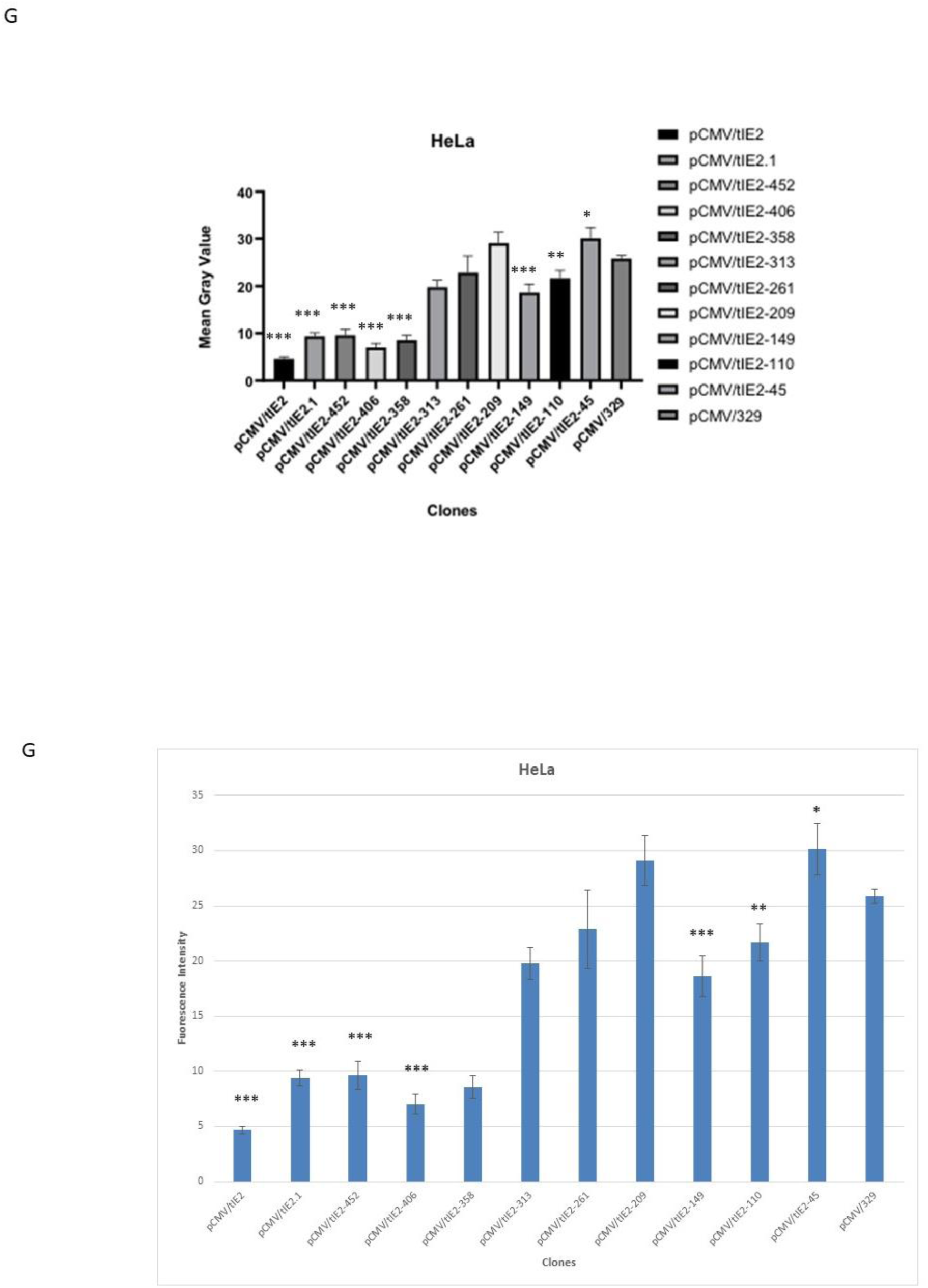
Identification of inhibitory sequences by 5’ end truncation analysis of the tIE2 promoter. (A) Diagrammatic representation of the OpIE2 promoter pointing out the different lengths of the truncated promoters. Each version was cloned separately in the pEGFP-N1 vector downstream CMV promoter as represented in figure 2, constructing different versions of the shuttle vector. (B) Fluorescence microscopy images of Sf9 cells after transfection with the nine versions of the shuttle vector containing nine different versions of the tIE2 promoter, and pCMV/tIE2 as a control. (C) A graph representing fluorescence intensity values for Sf9 cells in Figure 10B. (D) Fluorescence microscopy images in HEK-293T cells. (E) A graph representing fluorescence intensity values for HEK-293T cells in Figure 10D. (F) Fluorescence microscopy images of HeLa cells. (G) A graph representing fluorescence intensity values for HeLa cells in Figure 10F. All numerical values were obtained using ImageJ software. * represents statistical significance of P <0.05, ** represents significance of P<0.01, and *** represents significance of P< 0.001.

### Quantification of the expression level of pCMV/tIE-313, pCMV/tIE-261, and pCMV/tIE-209 vectors in Sf9 and HEK-293T cells

EGFP expression by pCMV/tIE-313, pCMV/tIE-261, and pCMV/tIE-209 clones was quantified in Sf9 and HEK-293T cells by RT-qPCR. In comparison to pCMV/tIE2ΔCMV as a control, pCMV/tIE2-313 expression level was the highest in Sf9 cells, followed by pCMV/tIE2-261, then pCMV/tIE2-209 (Figure 11A). On the contrary, in HEK-293T cells, pCMV/tIE2-209 expression was the highest, followed by pCMV/tIE2-261, then pCMV/tIE2-313, in comparison to pEGFP-N1 as a control (Figure 11B).

**Figure 11.**
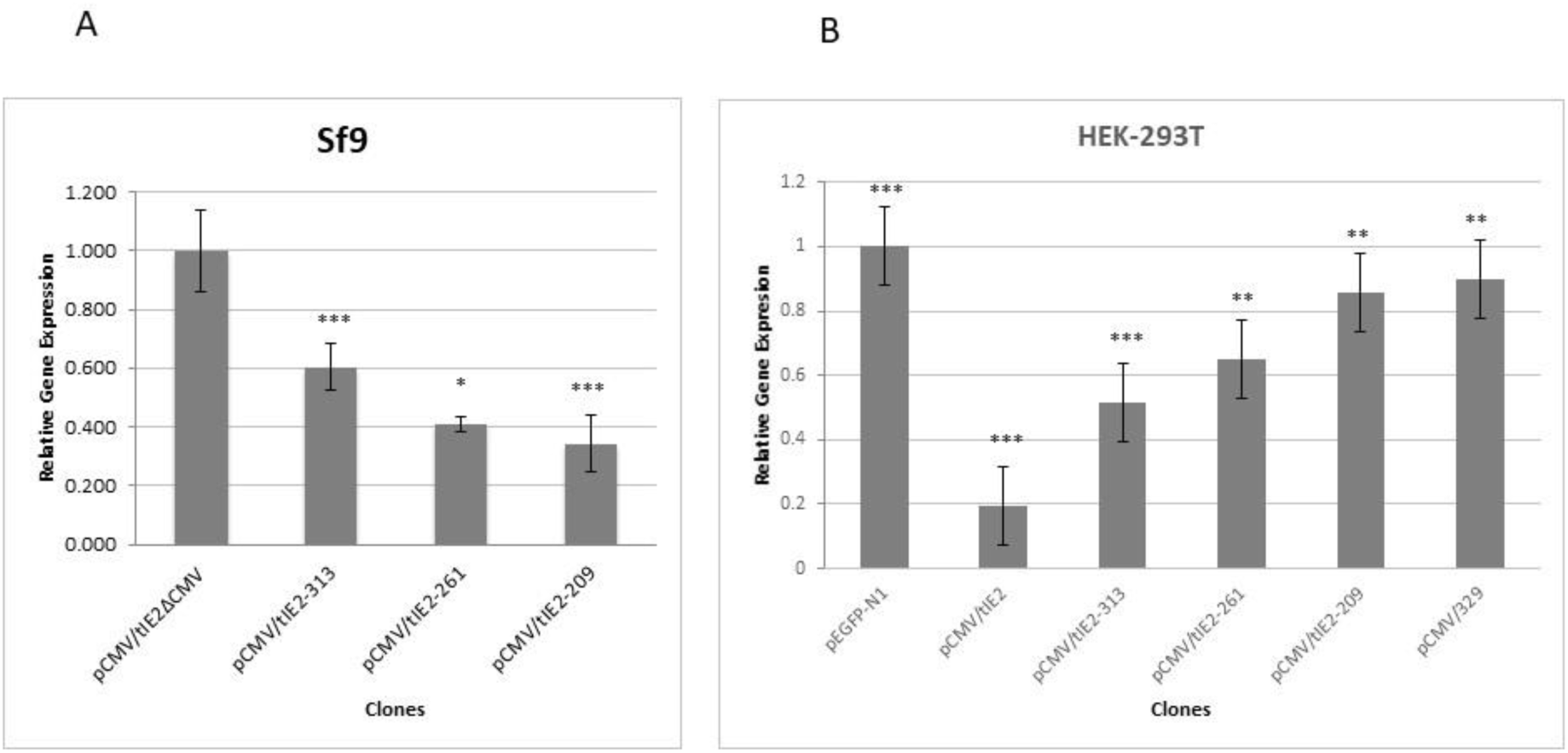
Quantification of EGFP transcript levels by qRT-PCR in transfected cells with plasmids pCMV/tIE2-313, pCMV/tIE2-261, and pCMV/tIE2-209. (A) EGFP transcript levels induced by pCMV/tIE2-313, pCMV/tIE2-261, pCMV/tIE2-209, and pCMV/tIE2ΔCMV vectors in Sf9 cells. (B) EGFP transcript levels induced by pEGFP-N1, pCMV/tIE2, pCMV/tIE2-313, pCMV/tIE2-261, pCMV/tIE2-209, and pCMV/329 vectors in HEK293T cells. These data values were obtained from the mean of three independent replicates. * represents statistical significance of P <0.05, ** represents significance of P<0.01, and *** represents significance of P< 0.001.

### FACS confirms the efficiency of the chimeric CMV/tIE2-261 promoter in Sf9 and HEK-293T cells

The chimeric CMV/tIE2-261 promoter’s activity was tested by flow cytometry in Sf9 and HEK-293T cells. The pCMV/tIE2ΔCMV and pEGFP-N1 vectors were used as controls. High expression levels were obtained by the shuttle vector in both cell lines (Figure 12).

**Figure 12.**
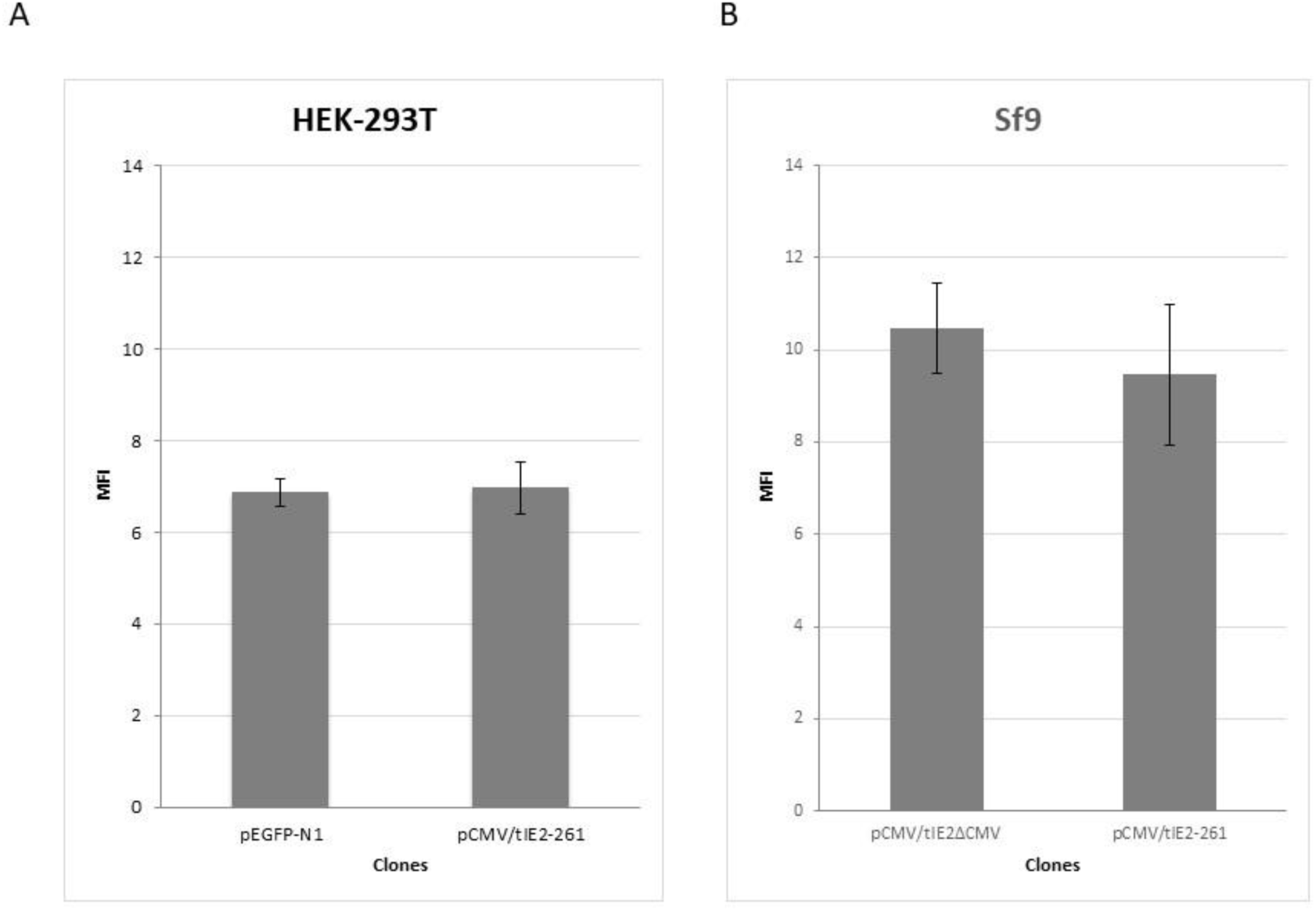
Measuring EGFP expression level in cells transfected with plasmid pCMV/tIE2-261 in HEK-293T and Sf9 cells by Flow Cytometry. (A) Mean fluorescence intensity for HEK-293T cells expressing EGFP by pCMV/tIE2-261 vector. pEGFP-N1 vector was used as a control. (B) Mean fluorescence intensity for Sf9 cells expressing EGFP by pCMV/tIE2-261 vector. The pCMV/tIE2ΔCMV vector was used as a control.

## DISCUSSION

The use of tandem promoters in eukaryotic cells is always challenging since many factors can directly affect the promoters’ activity including (I) transcriptional interference (Shearwin et al. 2005), (II) length of the proximal promoter sequence (Pande 2018), and (III) presence of ATGs in the downstream promoter (Philipps et al. 2008).

In this work, we studied the influence of CMV promoter on the OpIE2 promoter and vice versa in Sf9 insect cells, HEK-293T cells, and HeLa mammalian cells. The first challenge was the order of the promoters: which promoter should come first? The proposed order of the promoters was CMV followed by OpIE2 since the distance between the CMV promoter and the gene of interest was known not to be critical for its optimum activity (Philipps et al. 2008; Kozak 1999). The wild-type OpIE2 promoter sequence contains seven ATGs that might act as potential translational start sites, one of which is followed by a G at the +4 position that is an important element of the Kozak sequence in mammalian cells (Kozak 1989). A truncated version of the OpIE2 promoter was constructed to delete six ATG triplets, near the 5’ end of the promoter, out of a total of seven ATGs in the whole promoter sequence. Fortunately, those six deleted triplets lie in a region that is not known to be critical for the OpIE2 promoter activity in Sf9 cells, unlike the seventh ATG triplet (Theilmann and Stewart 1992). As this seventh ATG is located close to the TATA box of the promoter, modifying it may be critical to the promoter activity. Therefore, *in silico* analysis was performed to select the mutation that can eliminate this potential start codon with a minimal detrimental effect on the OpIE2 promoter activity in Sf9 cells. Although nine possible mutations can modify the ATG triplet (<u>T</u>TG, <u>C</u>TG, <u>G</u>TG, A<u>A</u>G, A<u>C</u>G, A<u>G</u>G, AT<u>C</u>, AT<u>T</u>, and AT<u>A</u>), the analysis did not detect any known regulatory protein that can bind to the ATG region even when all nine mutations were considered. The results suggested that mutating these three bases by any of the nine possibilities should not affect the binding of any transcription factor to the OpIE2 promoter in insect cells. This assumption was then tested experimentally.

Nine versions of −75 ATG-mutant OpIE2 promoter were successfully obtained and were efficiently functioning in Sf9 cells. However, the activity of the chimeric promoter (CMV/tIE2) was very poor in HEK-293T cells. This finding was supported by similar data obtained from different cell lines that are not shown (HeLa, MCF7, LnCap, and Huh7 cells). The in-gel fluorescence results implied that the decline in CMV promoter activity in mammalian cells was not due to the presence of ATG triplets in the OpIE2 promoter sequence. This suggested that the inhibition started at the transcription level, which was later confirmed by RT-qPCR. After dividing the truncated OpIE2 promoter sequence, and producing pCMV/tIE2.1 and pCMV/tIE2.2 clones, it was found that the region near the 5’ end (CMV/IE2.1) had a higher inhibitory effect on CMV promoter activity in mammalian cells than the region near the 3’ end (CMV/IE2.2). One explanation could be the possibility of transcriptional interference between the two promoters (Shearwin et al. 2005; Hao et al. 2017). Although OpIE2 is inactive in mammalian cell lines (Pfeifer et al. 1997), pairwise local alignment between the CMV promoter and the OpIE2 promoter indicated that the two promoters shared some sequence similarity. This suggests the possibility of imperfect TF recruitment on OpIE2 in mammalian cells. This insufficient or flawed recruitment is unable by itself to initiate the transcription process in mammalian cells, but at the same time can cause interference or collision with the RNA polymerase recruited on the CMV promoter (Turchinovich et al. 2016). This hypothesis was studied by transfecting HEK-293T cells with pCMV/329/IE2.1 and pCMV/329/IE2.2 vectors, which suggested that transcription interference does not contribute significantly to the low CMV promoter activity in these cells, or at least that it was not the main cause of reduction in CMV promoter activity.

5’ end consecutive deletions of the OpIE2 promoter were conducted to identify the inhibitory regions (Figure 10). By transfecting these vectors into mammalian and insect cells, it was noticed that as the OpIE2 fragment size shortened, the EGFP expression efficiency improved in mammalian cells and weakened in insect cells. This agrees with previous studies revealing the importance of an 18 bp element in the OpIE2 sequence repeated at the 3’ end of the promoter (Theilmann and Stewart 1992). This element was suggested to be composed of two smaller 8bp and 10bp elements (Theilmann and Stewart 1992). An enhancement in EGFP fluorescence levels in mammalian cells was observed starting from clone pCMV/tIE2-313. This suggested that the first 175 bp at the 5’ end of the tIE2 sequence specifically have an inhibitory effect on CMV promoter. Clones containing larger fragments of OpIE2 promoter sequence (pCMV/tIE-452, pCMV/tIE-406, pCMV/tIE-358) showed more inhibitory effect on the CMV promoter activity in mammalian cells. Nevertheless, this effect was linked to the above mentioned 175 pb inhibitory region rather than the OpIE2 promoter length and the distance it creates between the CMV promoter and the target gene. This assumption was based on the fact that clone pCMV/tIE2.1, which also contains these 175 bp in a shorter version of tIE2 promoter (329 bp), still had low CMV promoter activity in mammalian cells.

Although it was known in the literature that the OpIE2 promoter works efficiently up to −270 bp, shorter promoter fragments in pCMV/tIE2-261 and pCMV/tIE2-209 maintained high fluorescence levels in insect Sf9 cells (Theilmann and Stewart 1992). Clones pCMV/tIE-313, pCMV/tIE-261, pCMV/tIE-209 seemed to have acceptable EGFP fluorescence levels in Sf9 and HEK-293T cells, thus their expression levels were quantified using RT-qPCR in both cell lines. The three vectors showed opposite expression patterns in Sf9 and HEK-293T cell lines. As the length of the insect promoter OpIE2 decreases, its efficiency in the Sf9 cells decreases, and the CMV promoter activity in mammalian cells increases. This gradual change in CMV promoter activity in mammalian cells was thought-provoking since in terms of the distance created by the tIE2 promoter fragments between the CMV promoter and the target gene, the three tIE2 fragments (tIE2-313, tIE2-261, and tIE2-209) had acceptable lengths and should not affect the CMV promoter activity negatively. Also, judging by the high EGFP expression induced by the pCMV/329 vector, a 329 bp long space between the CMV promoter and the target gene was not an obstruction to the promoter’s activity. From this set of results, we concluded that the OpIE2 sequence itself has an inhibitory effect on CMV promoter activity. Furthermore, this inhibition, although concentrated at the 5’ end of the truncated OpIE2 promoter, is also scattered throughout the OpIE2 promoter sequence. We suggest that this inhibition is caused by multiple short sequences that may recruit or bind to proteins, which interfere with the transcription process halting the RNA polymerase motion.

This study establishes a platform that can be used to further understand how viral promoters work in non-host cells and how mammalian cells act against an exogenous regulatory sequence. Moreover, this extensive study of the baculovirus promoter (OpIE2) can give a better understanding of how this promoter works in different cell types. These results also recommend pCMV/tIE2-261 as an improved shuttle vector owing to its high expression level in insect and mammalian cell lines. Finally, the gradual inhibition associated with the different OpIE2 promoter lengths can serve to tailor the level of expression of the CMV promoter in mammalian cells by using a longer OpIE2 fragment downstream of the CMV promoter when a lower expression is needed and a shorter fragment when a higher expression is desired.

## ACKNOWLEDGEMENTS

The Authors thank Dr. Marwan Emara, director of the ‘‘Center of Aging and Associated Diseases’’ (CAAD) at Zewail City for his help and support throughout the work and Ms. Nahla Hassan for her help in reviewing the manuscript. Also, we would like to thank the ‘‘Genome Center’’ and the ‘‘Center of Excellence for Stem Cell Research’’ at Zewail City for allowing us to use their facilities during the initial study.

## Competing interests

The authors involved in this study declared that they do not have any conflict of interest from any kind related to this manuscript.

## FUNDING

This work is supported by the Science and Technology Development Fund (STDF) ID 22976 and in part with the internal fund ZC 016-2019.

## Supplementary Data

**Table S1:**
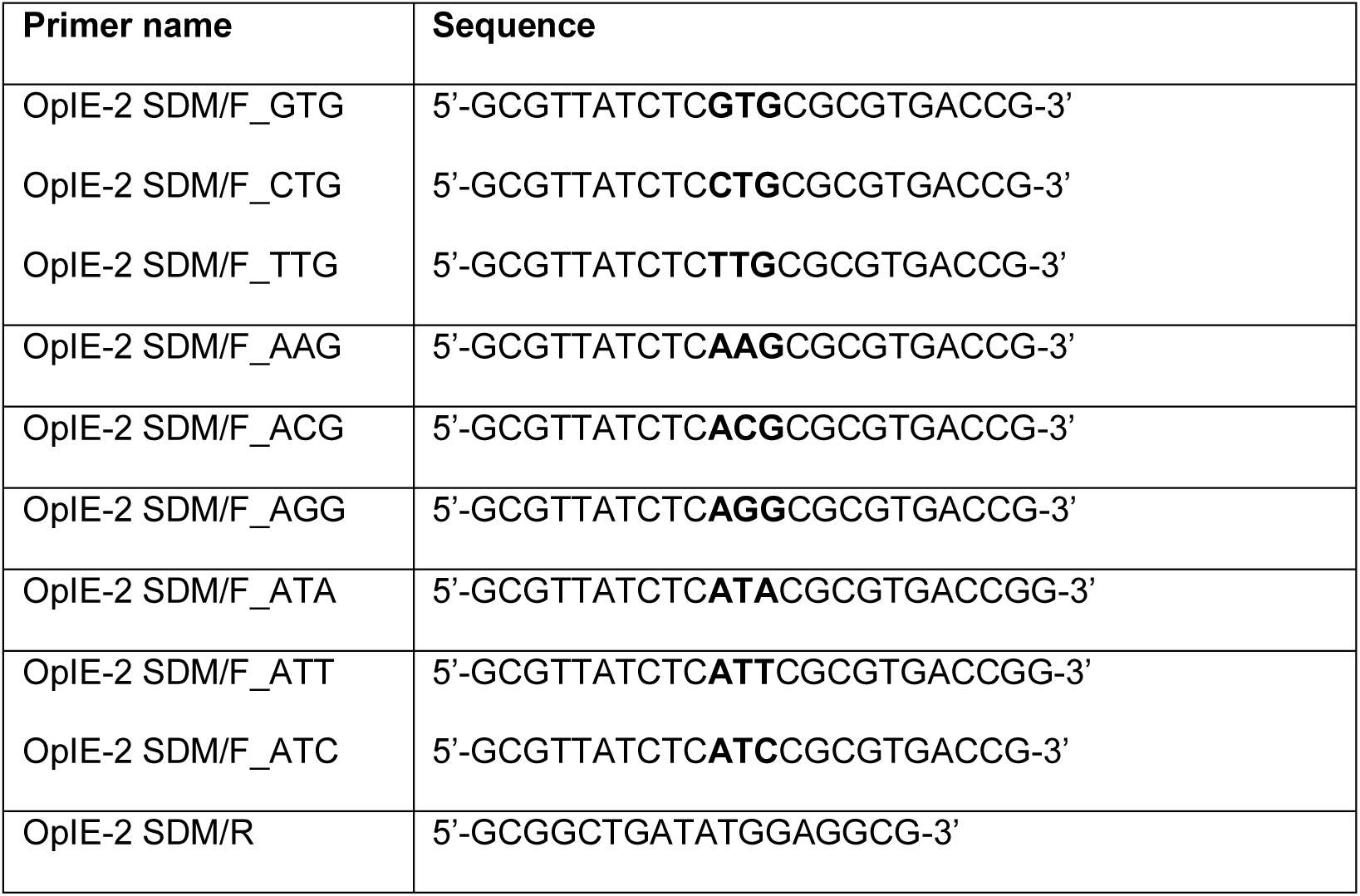
List of primers used for Site-directed mutagenesis to produce tIE2 promoter with no ATG triplet in its sequence.

**Table S2:**
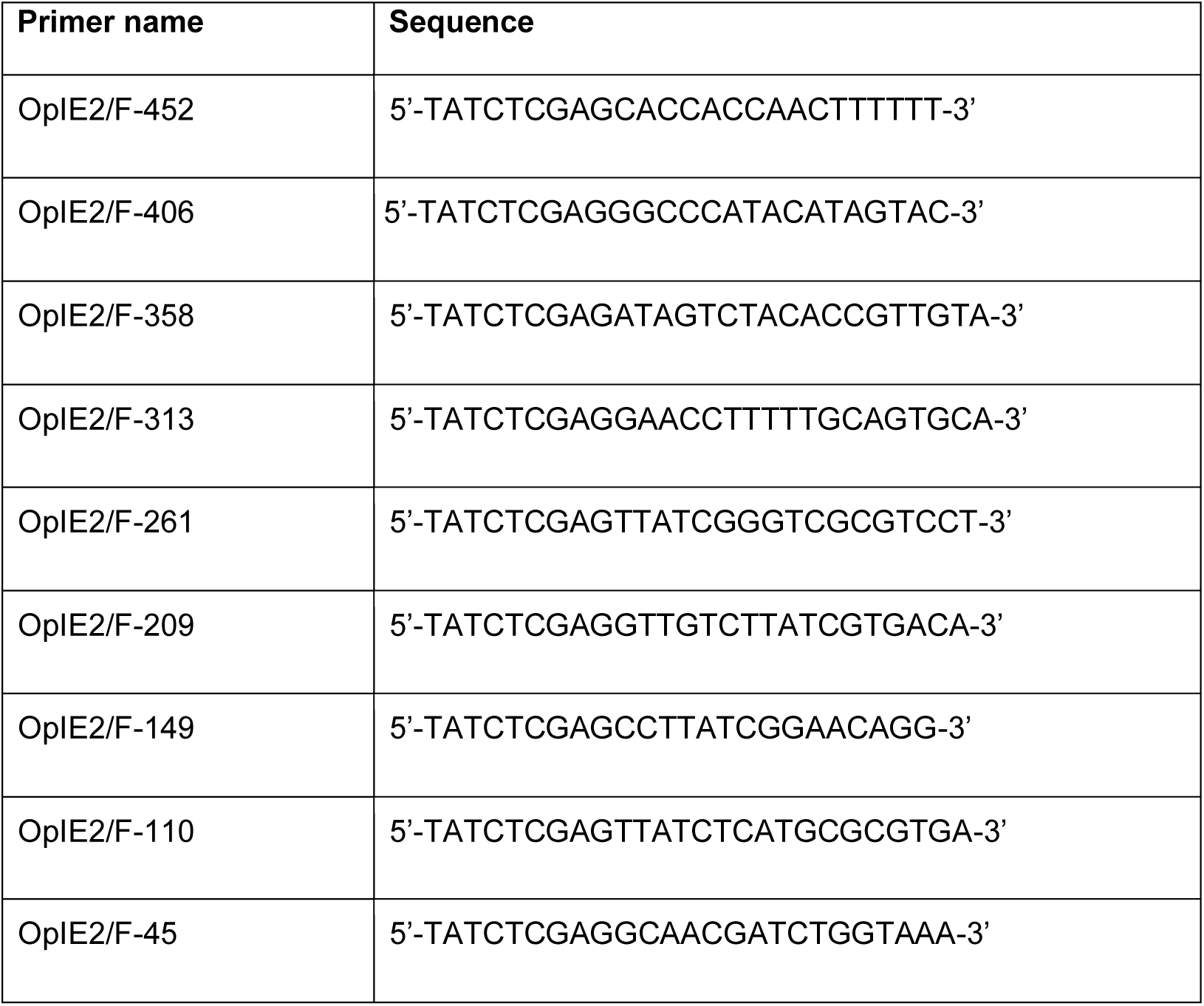
List of primers used for tIE2 truncation.

**Table S3:** Potential TFs that can bind to the repeated elements within the tIE2 promoter sequence.

| TF | Protein name | UniProt ID | Gene group/Protein Family | Consensus sequence | Annotated function |
| --- | --- | --- | --- | --- | --- |
| Achi | Achintya, isoform C | A1Z916 | TALE<br>Homeobox<br>TFs | TGACAG | DNA binding protein that activates the transcription (positive regulator) (Conlon et al. 1994). |
| Vis | Vismay, isoform B | A1Z913 | TALE<br>Homeobox<br>TFs | TGACAG | DNA binding protein that activates the transcription (positive regulator) (Hyman et al. 2003). |
| Hth | Homeobox protein homothorax | O46339 | TALE<br>Homeobox<br>TFs | TGACAG | Its annotated functions are related to the salivary glands eye development in *Drosophila melanogaster* (Henderson and Andrew 2000; Wernet et al. 2003). |
| GATAe | GATAe,<br>isoform A | Q9VF00 | GATA TFs | GCGATAAG | DNA binding protein that<br>activates the<br>transcription (positive<br>regulator) (Okumura et<br>al. 2007). |
| Optix | Protein Optix | Q95RW8 | SIX/Sine oculis<br>homeobox<br>family | TGATA | Transcriptional repressor<br>(Kenyon et al. 2005). |

## Notes

### Competing Interest Statement

The authors have declared no competing interest.

